# Salicylic Acid Engages Central Metabolic Regulators SnRK1 and TOR to Govern Immunity by Differential Phosphorylation of NPR1

**DOI:** 10.1101/2025.06.17.660129

**Authors:** Yixuan Chen, John Withers, Tianyuan Chen, Sargis Karapetyan, Jan Draken, Yezi Xiang, Wolfgang Dröge-Laser, Xinnian Dong

## Abstract

Immunity is a delicate balance between combating infection and preserving the metabolic functions vital for host survival. However, the mechanisms by which immune responses are coordinated with cellular metabolism remain largely unknown. Here, we show that NONEXPRESSER OF PR GENES 1 (NPR1), the central plant immune regulator of salicylic acid (SA)-mediated defense responses, is controlled by a cascade of posttranslational modifications (PTMs) involving two master nutrient-sensing kinases. Under normal growth conditions, TARGET OF RAPAMYCIN (TOR) inhibits NPR1 through phosphorylation at Ser-55/59. During defense responses, elevated SA enhances SNF1-RELATED KINASE 1 (SnRK1) activity, which in turn inhibits TOR signaling and phosphorylates NPR1 at Ser-557. This phosphorylation event activates NPR1 and facilitates its subsequent PTMs. Together, our results reveal an integral role of SA (the active metabolite of aspirin) in controlling central metabolic regulators SnRK1 and TOR to coordinate immune responses and growth through antagonistic modifications of NPR1.

## INTRODUCTION

Plants have evolved a sophisticated innate immune system that can be activated by plasma membrane-associated pattern recognition receptors upon detection of microbe-associated molecular patterns^1,2^ and by intracellular nucleotide-binding leucine-rich repeat immune receptors which respond to specific pathogen effectors^3^. The resulting pattern-triggered immunity (PTI) and effector triggered immunity (ETI) are defense mechanisms to mitigate local infection^4^. Interestingly, the local immune response can also lead to the establishment of a long-lasting, broad-spectrum defense response known as systemic acquired resistance (SAR) mediated by the immune hormone salicylic acid (SA)^5-7^. An increase in endogenous SA synthesis triggered by infection or exogenous application of SA induces massive transcriptional reprogramming, leading to the production and secretion of antimicrobial proteins, such as PATHOGENESIS-RELATED PROTEIN 1 (PR1)^8-10^ and the establishment of SAR. This SA-mediated immune response requires the function of NONEXPRESSER OF PR GENES 1 (NPR1) which was first identified through genetic screens in *Arabidopsis*^8,9^ and found to have single-copy orthologs in most of the angiosperm species except *Brassicacea*^11^. NPR1 is required for initiating the SA-mediated transcriptional cascade as a cofactor for the TGA group of transcription factors (TFs)^12-14^.

Though the NPR1 protein is constitutively expressed, its activity and stability are tightly controlled through post-translational modifications (PTMs)^15^. In its uninduced state, NPR1 is phosphorylated at Ser-55 and Ser-59 (Ser-55/59)^16^ and is sequestered in the cytoplasm as an oligomer via disulfide bond formation^17^. Upon SA induction, NPR1 is reduced by thioredoxins to antagonize the effect of S-nitrosylation^18^ and undergoes dephosphorylation at Ser-55/59^16^. These modifications enable NPR1 to translocate into the nucleus. In the nucleus, NPR1 is SUMOylated by the SMALL UBIQUITIN-LIKE MODIFIER 3 (SUMO3)^16^ and phosphorylated at Ser-11 and Ser-15 (Ser-11/15)^19^ to promote its interaction with the ubiquitin/proteasome system, leading to its full activation and subsequent degradation^19,20^. Disruption of these PTMs has been shown to result in either defects in SAR or autoimmunity^16,19,20^. However, few kinases have previously been implicated in regulating NPR1. PKS5/SnRK3.22 phosphorylates NPR1 at the C-terminal region, altering its interaction with WRKY TFs^21^, while SnRK2.8, a key abscisic acid (ABA) signaling component, phosphorylates NPR1 at Thr-373 and Ser-589^22^. However, the dynamic regulation of enzymes that modify NPR1 at different sites in response to SA remains underexplored.

Despite being primarily studied as a defense hormone, SA has previously been shown to affect plant metabolism. SA production has been linked to alternative respiration via alternative oxidase (AOX) during thermogenesis^23,24^, and AOX also contributes to stress resilience by maintaining redox and metabolic homeostasis^25^. SA has also been demonstrated to bind to CATALASE 2, inhibiting its scavenging of hydrogen peroxide^26,27^, while the redox changes resulting from SA treatment induce the nuclear translocation of NPR1^17^. However, how metabolic pathways influence the SA-mediated immune response remains a significant knowledge gap.

Like in other eukaryotes, plant metabolic activities are controlled by the two central nutrient-sensing kinases: TARGET OF RAPAMYCIN (TOR)^28-30^ and SNF1-RELATED PROTEIN KINASE 1 (SnRK1)^31-33^, also known as SUCROSE NON-FERMENTING 1/AMP-ACTIVATED PROTEIN KINASE (SNF1/AMPK) in yeast and mammals. TOR is activated in nutrient-rich conditions, driving anabolic and energy-demanding processes and promoting growth by activating translation-related proteins^30,34,35^. Conversely, SnRK1 senses various abiotic stresses, such as prolonged darkness, exposure to herbicides, hypoxia, and treatment by the stress hormone abscisic acid (ABA)^32,36,37^. When activated by abiotic stresses, SnRK1 suppresses TOR activity to inhibit growth and causes a shift towards stress tolerance^38^. Though SnRK1 and TOR have also been implicated in defense responses against pathogens^38-46^, how these kinases are mechanistically linked to plant immunity is poorly understood.

Here, we identified NPR1 as a direct phosphorylation substrate of SnRK1 and TOR. SA activates SnRK1 to phosphorylate NPR1 at Ser-557, a new PTM essential for NPR1 function. Furthermore, SA-activated SnRK1 suppresses the TOR activity, blocking its phosphorylation of NPR1 at Ser-55/59, thereby facilitating NPR1’s release and activation from its quiescent state. Therefore, our study reveals a direct link between SA-mediated immunity and cellular metabolic pathways through the differential phosphorylation of NPR1 by SnRK1 and TOR.

## RESULTS

### Ser-557 phosphorylation is required for SA-mediated activation of NPR1

To identify novel PTM sites in NPR1 beyond those discovered through targeted approaches based on known phosphorylation consensus^16,19^, we employed proteomics to detect NPR1-specific phospho-peptides from mature *Arabidopsis* after SA treatment. Targeted mass spectrometry revealed a previously undiscovered phosphorylation at Ser-557 in NPR1 peptides following proteolytic digestion with two different enzymes, trypsin and GluC (Figure 1A). To confirm this result, we developed a phospho-specific antibody against Ser-557 and found that this modification occurs in transgenic *Arabidopsis* (*35S:NPR1-GFP/npr1-2*) plants following SA treatment and in response to infection by the bacterial pathogen *Pseudomonas syringae* pv. *maculicola* ES4326 (*Psm* ES4326) (top, Figure 1B). The endogenous NPR1 is similarly phosphorylated in wildtype Col-0 (WT) plants upon SA treatment (bottom, Figure 1B).

**Figure 1.**
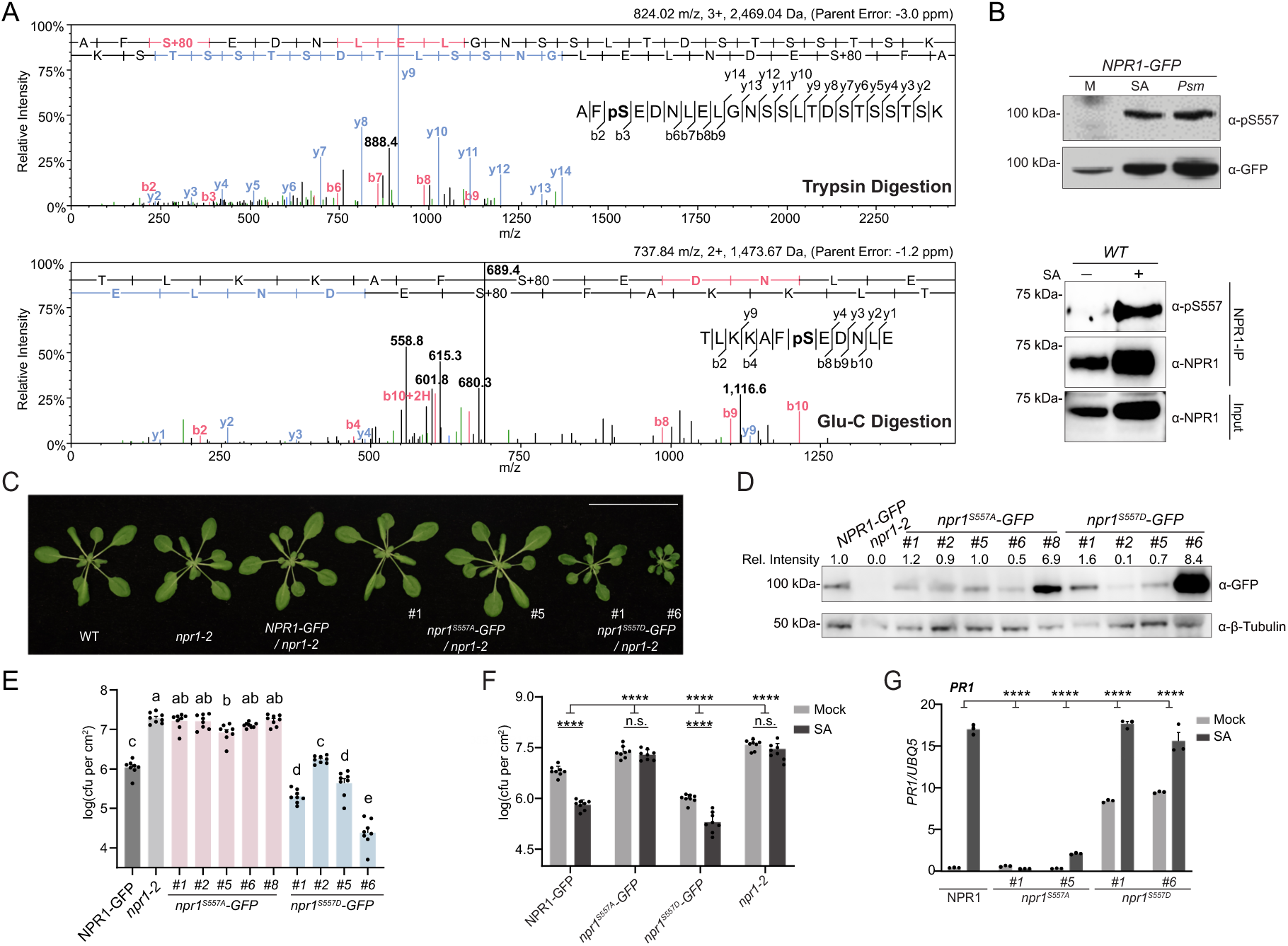
Phosphorylation of NPR1 at Ser-557 is necessary and partially sufficient for SA-induced disease resistance. (A)Liquid chromatography-tandem mass spectrometry (LC-MS/MS) analysis of NPR1 phosphorylation. *35S:NPR1-GFP* (in *npr1-2*) plants were treated with mock or 0.5 mM SA for 24 h, followed by trypsin (top)/Glu-C (bottom) digestion for LC-MS/MS. (B)Phosphorylation of NPR1 at Ser-557. Top: *35S:NPR1-GFP* plants were treated with mock (M), 0.5 mM SA (SA) for 24 h or inoculated with *Pseudomonas syringae* pv. *maculicola* ES4326 (*Psm*, OD_600_ = 0.0002). Bottom: Wild-type Col-0 (WT) plants were treated with mock (-) or 1 mM SA (+) for 8 h. The endogenous NPR1 protein was immunoprecipitated using protein A/G magnetic agarose conjugated to the α-NPR1 antibody. Samples were western blotted with an antibody specific to phosphorylated Ser-557 (α-pS557). Total NPR1 protein levels were verified using the α-GFP and α-NPR1 antibody, respectively. (C)Representative photographs of 3.5-week-old WT, *npr1-2*, *35S:NPR1-GFP*, and independent *35S:npr1^S557A^-GFP* (in *npr1-2*) and *35S:npr1^S557D^-GFP* (in *npr1-2*) transgenic lines (#number). Scale bar, 5 cm. (D) NPR1-GFP levels in *35S:NPR1-GFP* and in independent transgenic *35S:npr1^S557A^-GFP* and *35S:npr1^S557D^-GFP* lines (#numbers) were determined using the α-GFP antibody. β-tubulin served as the loading control to calculate relative (rel.) intensity of the NPR1-GFP bands. (E) Basal resistance in *35S:NPR1-GFP*, *35S:npr1^S557A^-GFP, 35S:npr1^S557D^-GFP*, and *npr1-2* plants inoculated with *Psm* ES4326 (OD_600_ = 0.0002). Bacterial growth was measured at 3 days post-inoculation (dpi). cfu, colony-forming units. *n* = 8. (F) SA-mediated resistance in *35S:NPR1-GFP*, *35S:npr1^S557A^-GFP #1*, *35S:npr1^S557D^-GFP #1*, and *npr1-2* plants. 3.5-week-old plants were treated with mock or 1 mM SA, followed by inoculation with *Psm* ES4326 (OD_600_ = 0.001) 24 h later. Bacterial growth was measured at 3 dpi. *n* = 8. (G) *PR1* gene expression. 3.5-week-old *35S:NPR1-GFP*, *35S:npr1^S557A^-GFP*, and *35S:npr1^S557D^-GFP* plants treated with mock or 1 mM SA for 24 h. *PR1* transcript levels were determined by RT-qPCR, and *UBQ5* expression was used for normalization. All data are presented as mean ± s.e.m. Individual columns were compared using one-way ANOVA with Tukey’s post-hoc (E) or a two-tailed Student’s t-test (F), and the interactions were tested using two-way ANOVA (F and G), ****p < 0.0001. n.s., not significant; different lowercase letters indicate statistical significance tested between multiple groups by one-way ANOVA at p < 0.05. See also Figure S1.

**Figure S1.**
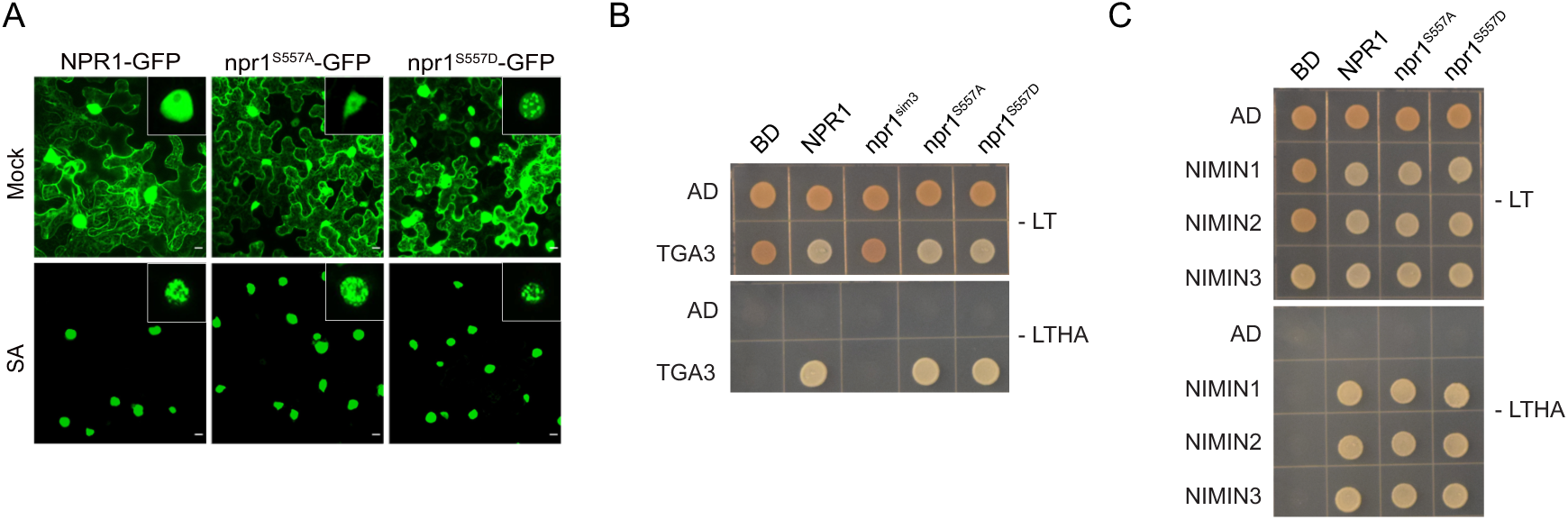
Ser-557 phosphorylation facilitates nSINC formation without affecting NPR1 localization or its interactions with TGA3 and NIMIN1-3, related to. **Figure 1**. (A) NPR1 subcellular localization. Confocal micrographs of *N. benthamiana* leaves transiently expressing *35S:NPR1-GFP*, *35S:npr1^S557A^-GFP* or *35S:npr1^S557D^-GFP* at 30 h post-Agrobacterium infiltration. Mock or 1 mM SA treatments were applied 24 h post-infiltration. Insets highlight nuclear condensates. Scale bars, 10 μm. (B and C) Yeast two-hybrid interaction between BD-NPR1 (NPR1), BD-npr1^S557A^ (S557A), or BD-npr1^S557D^ (S557D) with AD-TGA3 (TGA3) (B) or AD-NIMIN1 (NIMIN1), AD-NIMIN2 (NIMIN2), and AD-NIMIN3 (NIMIN3) (C). –LT, synthetic dropout media lacking Leu and Trp; –LTHA, media lacking Leu, Trp, His, and Ade.

To investigate whether phosphorylation of NPR1 at Ser-557 contributes to its function, we generated phospho-dead *35S:npr1^S557A^-GFP* (*npr1^S557A^-GFP*) and phospho-mimetic *35S:npr1^S557D^-GFP* (*npr1^S557D^-GFP*) mutant lines in the *npr1-2* background. While no obvious growth phenotype was observed in the *npr1^S557A^-GFP* plants, the *npr1^S557D^-GFP* lines exhibited varying degrees of stunted growth that correlated with npr1^S557D^ protein levels (Figures 1C and 1D). Hence, we hypothesized that the phospho-mimetic mutant of Ser-557 might represent an auto-active or partially active form of NPR1. To test our hypothesis, we examined the basal resistance of *NPR1-GFP*, *npr1^S557A^-GFP*, and *npr1^S557D^-GFP* plants in response to *Psm* ES4326 (OD_600_ = 0.0002) and found that, similar to *npr1-2*, the *npr1^S557A^-GFP* plants had a higher than *NPR1-GFP* levels of bacterial growth, regardless of the npr1^S557A^-GFP protein levels (Figures 1D and 1E). In contrast, different *npr1^S557D^-GFP* lines showed varying levels of bacterial growth, with lines expressing higher protein levels showing less *Psm* ES4326 growth. We then tested the mutant plants for SA-induced resistance using a high inoculant of *Psm* ES4326 (OD_600_ = 0.001). We found that *npr1^S557A^-GFP* plants were insensitive to SA treatment while the *npr1^S557D^-GFP* plants exhibited higher resistance under both basal and SA-treated conditions compared to *NPR1-GFP*, but had less pronounced sensitivity to SA-induced resistance (Figure 1F).

To understand how Ser-557 phosphorylation contributes to SA- and NPR1-mediated immunity at the cellular and molecular level, we first tested whether the phosphorylation alters NPR1’s nuclear translocation in response to SA. We found that neither npr1^S557A^ nor npr1^S557D^ mutation had a significant effect (Figure S1A). Interestingly, even in the absence of SA treatment, npr1^S557D^-GFP formed nuclear biomolecular condensates (nSINC), which are normally observed after SA treatment in NPR1-GFP. We next performed yeast two-hybrid assays to examine the effects of Ser-557 phosphorylation on NPR1’s interaction with TGA3 TF, whose transcriptional activity is directly regulated by NPR1^13,14^. However, we detected no defect in the interaction, unlike npr1^sim3^, a mutant defective in SUMOylation by SUMO3^16^ (Figure S1B). Additionally, both Ser-557 phosphorylation variants were able to interact with NIMINs, which are known negative regulators of SAR that interact with NPR1^47,48^ (Figure S1C). Since Ser-557 phosphorylation does not appear to affect NPR1 nuclear translocation or its interaction with target TFs, we next examined the impact of this PTM on NPR1-mediated transcriptional activation, using *PR1* as a marker. We found that SA-induced expression of *PR1* was abolished in *npr1^S557A^-GFP* lines, while *npr1^S557D^-GFP* plants showed higher basal *PR1* levels, which were further enhanced by SA treatment (Figure 1G). These results further support our hypothesis that phosphorylation of Ser-557 is necessary, but only partially sufficient for NPR1 activation by SA.

### Elucidating the PTM cascade in SA-mediated activation of NPR1

To map the spatial and temporal dynamics of NPR1 modifications, we first investigated the subcellular compartment where Ser-557 phosphorylation occurs. We employed the *35S:NPR1-HBD* transgenic line in which SA-induced NPR1 nuclear translocation is blocked by the rat glucocorticoid receptor (GR) hormone-binding domain (HBD) due to its association with the heat shock protein HSP90 in the cytoplasm^49^. We found that releasing this association by application of the glucocorticoid hormone dexamethasone (DEX), in parallel with SA treatment, led to a significant increase in Ser-557 phosphorylation compared to the control (i.e., SA without DEX), indicating that this PTM predominantly occurs in the nucleus (Figure 2A).

**Figure 2.**
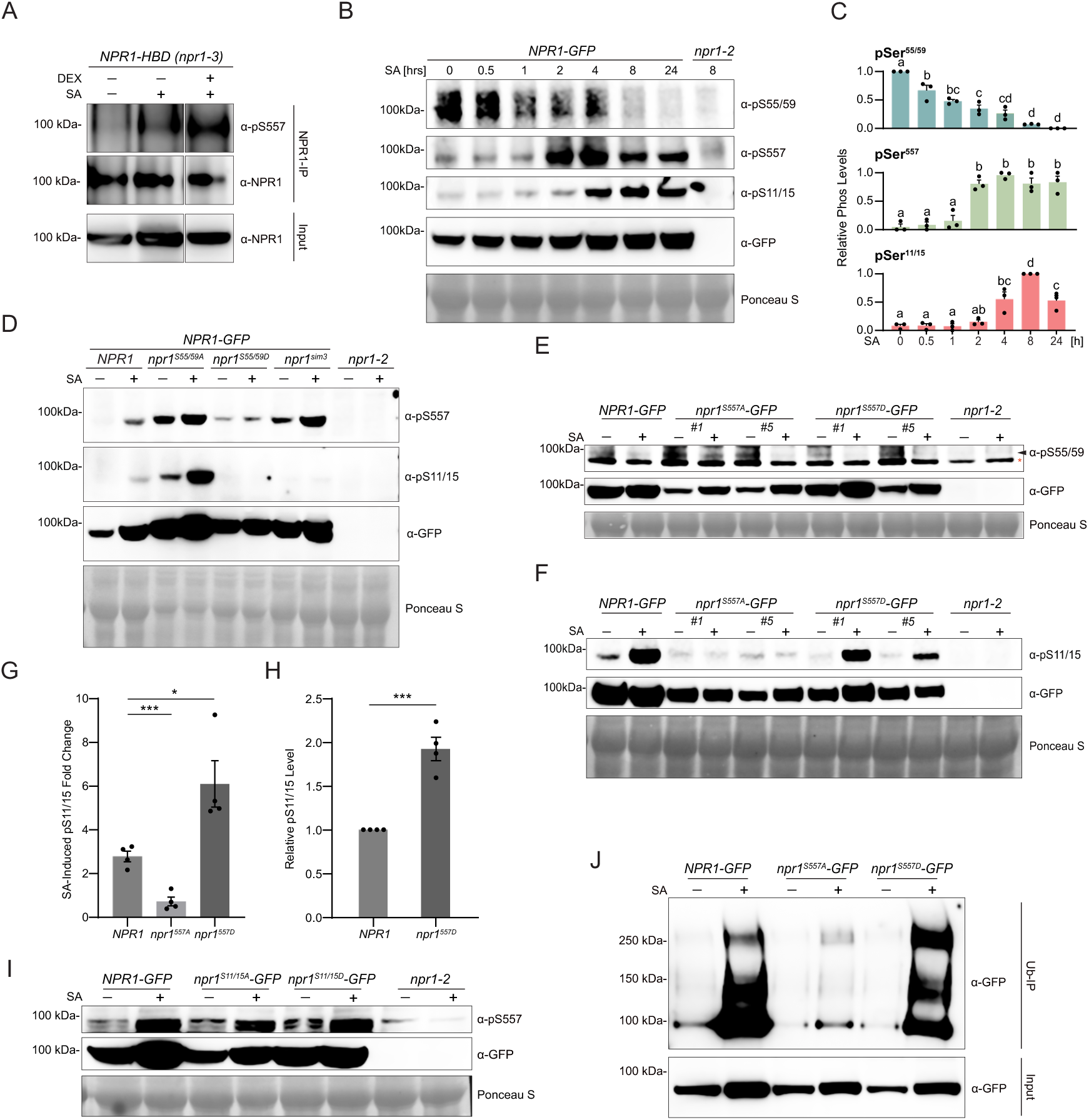
SA activates NPR1 through sequential post-translational modifications. (A) Subcellular location of NPR1 Ser-557 phosphorylation. *35S:NPR1-HBD* (in *npr1-3*) seedlings were treated with mock, 0.3 mM SA or 0.3 mM SA + 5 µM dexamethasone (Dex) for 8 h. NPR1-HBD protein was immunoprecipitated using protein A/G magnetic agarose coupled with the α-NPR1 antibody. Total and immunoprecipitated proteins were analyzed by western blotting with the indicated antibodies. (B and C) Time course of SA-induced NPR1 phosphorylation. *35S:NPR1-GFP* and *npr1-2* seedlings were treated with 0.3 mM SA for the indicated durations. Total protein was analyzed by western blotting. NPR1 phosphorylation was detected using antibodies against phosphorylated Ser-55 and Ser-59 (α-pS55/59), Ser-11 and Ser-15 (α-pS11/15), and Ser-557 (α-pS557) (B) and quantify in (C). NPR1 protein levels were verified using the α-GFP antibody and equal loading was confirmed using Ponceau S staining. For each blot, the band with the highest relative phosphorylation level (normalized to NPR1 protein levels) was set to 1. *n* = 3. (D) Hierarchy in NPR1 PTMs. *35S:NPR1-GFP*, *35S:npr1^S55/59A^-GFP* (in *npr1-2*), *35S:npr1^S55/59D^-GFP* (in *npr1-2*), *35S:npr1^sim3^-GFP* (in *npr1-2*), and *npr1-2* seedlings were treated as in (B) for 6 h. Total protein was analyzed by western blotting using α-pS557, α-pS11/15, and α-GFP antibodies. (E and F) Analysis of Ser-55/59 and Ser-11/15 phosphorylation in Ser-557 phospho-mutant lines. *35S:NPR1-GFP*, and independent *35S:npr1^S557A^-GFP*, *35S:npr1^S557D^-GFP* transgenic lines (#number), and *npr1-2* seedlings were treated as in (D). Total protein was analyzed with α-pS55/59 (E), α-pS11/15 (F), and α-GFP antibodies. *Non-specific band. (G and H) Quantification of Ser-11/15 phosphorylation in Ser-557 mutants from independent experiments. SA-induced fold changes in Ser-11/15 phosphorylation of NPR1-GFP, npr1^S557A^-GFP, and npr1^S557D^-GFP relative to mock (G). SA-induced Ser-11/15 phosphorylation in npr1^S557D^-GFP relative to that observed in NPR1-GFP after SA treatment (H). *n* = 4. (I) Analysis of Ser-557 phosphorylation in Ser-11/15 phospho-mutant lines. *35S:NPR1-GFP*, *35S:npr1^S11/15A^-GFP* (in *npr1-2*), *35S:npr1^S11/15D^-GFP* (in *npr1-2*), and *npr1-2* seedlings were treated and analyzed as in (D) using the α-pS557 and α-GFP antibodies. (J) Analysis of NPR1 ubiquitination in Ser-557 phospho-mutant lines. *35S:NPR1-GFP*, *35S:npr1^S557A^-GFP*, and *35S:npr1^S557D^-GFP* seedlings were treated as in (D). NPR1-GFP protein was immunoprecipitated using the Ubiquitin-trap. Total and immunoprecipitated proteins were analyzed with the α-GFP antibody. All data are presented as mean ± s.e.m. Individual columns were compared using one-way ANOVA with Tukey’s post-hoc (C) or a two-tailed Student’s t-test (G and H), *p < 0.05, ***p < 0.001. n.s., not significant; different lowercase letters indicate statistical significance tested between multiple groups by one-way ANOVA at p < 0.05.

Next, we sought to uncover the temporal relationships among the NPR1 phosphorylation sites known to affect plant defense. Phosphorylation at Ser-55/59 suppresses NPR1 activity^16,50^, while Ser-11/15 phosphorylation promotes immunity^19^. We performed a time course experiment to monitor changes in phosphorylation levels of Ser-55/59, Ser-557, and Ser-11/15 following SA treatment, using phospho-specific antibodies against each of these specific residues. We found that Ser-55/59 dephosphorylation occurred within 30 minutes of SA treatment and continued to decrease at later time points (Figures 2B and 2C). Ser-557 phosphorylation followed the dephosphorylation of Ser-55/59, occurring as early as two hours post-treatment. Ser-11/15 phosphorylation was induced later, at four hours post-SA treatment, and peaked at approximately eight hours. These results suggest a cascade of PTMs during SA-induced activation of NPR1.

We next used genetic approaches to investigate the sequential dependency of PTMs in activating NPR1. We previously found that npr1^S55/59A^ exhibited an auto-immune phenotype and caused Ser-11/15 phosphorylation even without SA treatment, while npr1^S55/59D^ failed to induce either Ser-11/15 phosphorylation or SAR^16^. We observed that npr1^S55/59A^ triggered Ser-557 phosphorylation under mock conditions, which was further enhanced by SA treatment, while npr1^S55/59D^ failed to induce Ser-557 phosphorylation even after SA treatment (Figure 2D). Conversely, neither npr1^S557A^ nor npr1^S557D^ mutation affected Ser-55/59 dephosphorylation (Figure 2E). These results indicate that Ser-557 modification occurs downstream of and is dependent on Ser-55/59 dephosphorylation. We also found that while the npr1^sim3^ SUMOylation mutant completely inhibited Ser-11/15 phosphorylation, it had no significant effect on Ser-557 phosphorylation, indicating that NPR1 SUMOylation occurs either downstream of or independent of Ser-557 modification. Ser-11/15 phosphorylation, which peaks later than Ser-557 phosphorylation, was blocked by npr1^S557A^ but enhanced by npr1^S557D^ after SA treatment (Figures 2F-2H), indicating that Ser-11/15 phosphorylation is dependent on Ser-557 phosphorylation. This result was further confirmed by the normal Ser-557 phosphorylation in both Ser-11/15 phospho-dead and phospho-mimetic mutants (Figure 2I). However, Ser-11/15 is not constitutively phosphorylated in *npr1^S557D^-GFP* under mock conditions, indicating that while Ser-557 phosphorylation is necessary, it is not sufficient for Ser-11/15 modification, a downstream PTM likely mediated by a currently unknown SA-dependent kinase. Finally, consistent with Ser-11/15 phosphorylation being required for NPR1 ubiquitination involved in its activation and degradation, a progressive event mediated by Cullin-RING E3 ligase and the E4 ligase UBE4^20^, we found that npr1^S557D^ could be ubiquitinated like NPR1, while npr1^S557A^ showed diminished ubiquitination, likely due to the lack of Ser-11/15 phosphorylation (Figure 2J).

### SnRK1 directly phosphorylates NPR1 at Ser-557

To identify the kinase that is responsible for Ser-557 phosphorylation, we focused on the SnRK family of kinases as they are known for their roles in stress responses^51-54^, and, more importantly, the Ser-557 site matches with the consensus sequence for SnRK1, Ф-x-Basic-xx-Ser-xxx-Ф (where x is any residue and Ф is a hydrophobic residue)^55,56^. We screened multiple SnRK family kinases expressed in *N. benthamiana* by performing *in vitro* phosphorylation assays using the Ser-557 phospho-antibody (α-pS557). We found that SnRK1α2, a catalytic subunit of SnRK1, could phosphorylate NPR1 at Ser-557 and this modification can be removed by lambda phosphatase (Figure S2A). Since *Arabidopsis* expresses the SnRK1α1 and SnRK1α2 subunits, we further tested them individually *in vitro*. We found that *in vitro* synthesized SnRK1α1 and SnRK1α2 could phosphorylate Ser-557 at low levels, consistent with their low catalytic activity, as indicated by moderate phosphorylation of the conserved T-loop (Thr-175 in SnRK1α1 and Thr-176 in SnRK1α2)^57,58^ detected using an α-pAMPK T-loop phospho-specific antibody (Figure 3A). Furthermore, Ser-557 phosphorylation was significantly enhanced by the addition of the SnRK1-ACTIVATING KINASE 1 (SnAK1), which by itself could not phosphorylate Ser-557, confirming that SnRK1 is the kinase phosphorylating Ser-557 *in vitro*. We also assessed the overall phosphorylation of NPR1 using a general phospho-serine/threonine antibody and found diminished phosphorylation in the npr1^S557A^ mutant (Figure 3A). Mass spectrometry analysis confirmed that only Ser-557 phosphorylation was increased with activated SnRK1, while other detectable phosphorylated serine or threonine residues showed minimal changes, indicating that SnRK1 specifically targets Ser-557 (Figures 3B and S2B). Co-immunoprecipitation (Co-IP) assays performed in *N. benthamiana* and transgenic *Arabidopsis* lines revealed that both SnRK1α subunits could interact with NPR1 *in vivo* (Figures 3C) and the interaction was further enhanced upon SA treatment (Figure 3D), likely due to increased NPR1 nuclear translocation. Confirming this hypothesis, pull-down assays indicated that SnRK1α1 and SnRK1α2 interact with NPR1 *in vitro* independent of their phosphorylation status by SnAK1 (Figure 3E). To confirm that SnRK1 phosphorylates Ser-557 *in vivo*, we mock-or SA-treated NPR1-GFP-expressing protoplasts from WT plants and two independent *snrk1α1^−/−^ snrk1α2^+/−^* (*sesquiα2*) mutants since double *snrk1α1 snrk1α2* knockout mutant is lethal^59,60^. We found that both *sesquiα2* lines were defective in SA-induced Ser-557 phosphorylation, indicating that SnRK1 is required for the phosphorylation of Ser-557 *in vivo* (Figure 3F). We also observed a defect in SA-mediated Ser-557 phosphorylation in mature *sesquiα2-1* plants (Figure 3G), further confirming our results *in vivo*. Combining both *in vitro* and *in planta* data, we concluded that SnRK1 is the *bona fide* kinase of Ser-557 in *Arabidopsis*.

**Figure 3.**
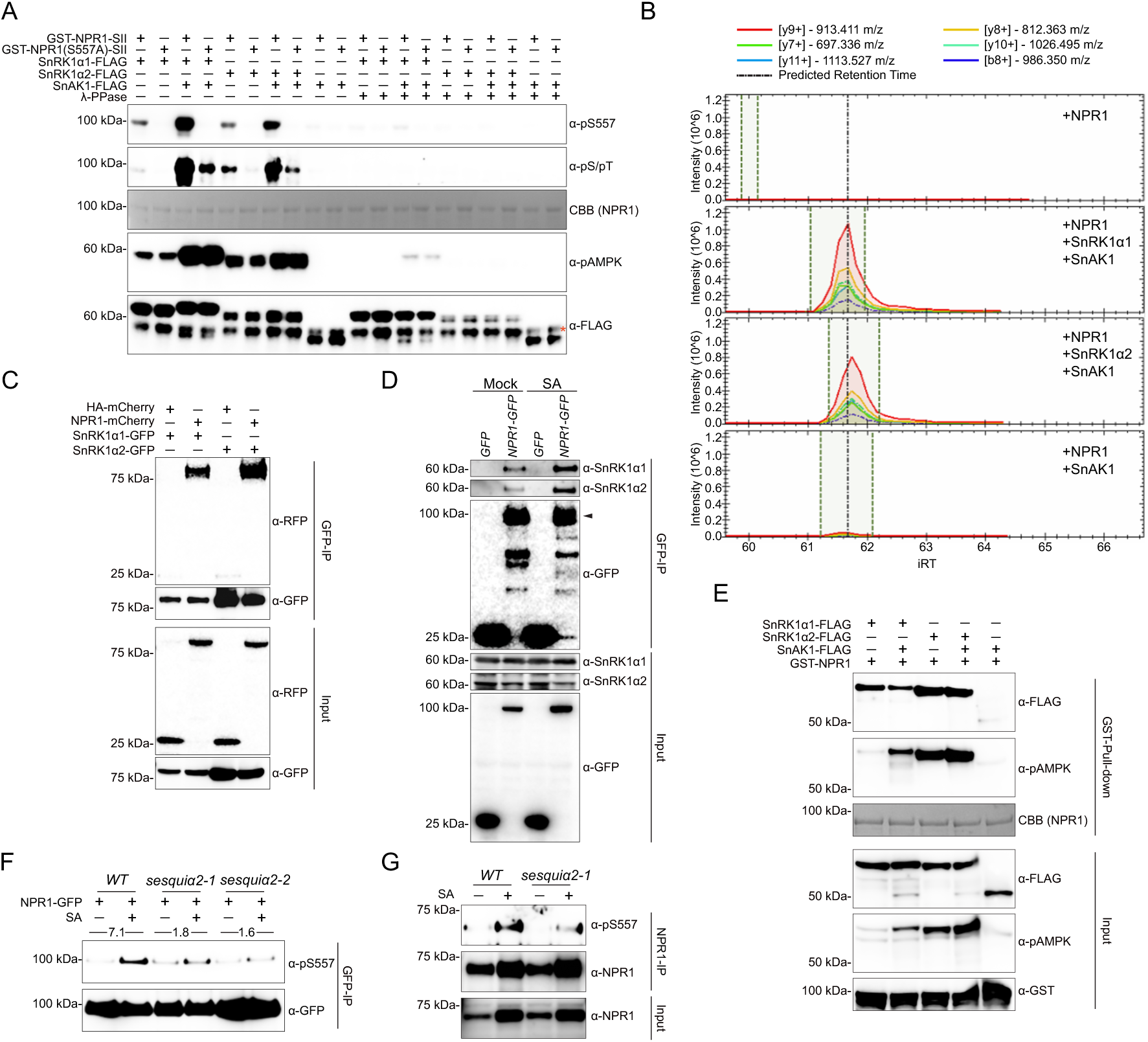
SnRK1 phosphorylates NPR1 at Ser-557 in response to SA induction. (A) *In vitro* phosphorylation assays of recombinant NPR1. Ser-557 phosphorylation was detected using the α-pS557 antibody and the overall phosphorylation of NPR1 was detected using the α-phospho-Ser/Thr antibody (α-pS/pT). SnRK1 activity was estimated by measuring phosphorylation of its T-loop using the α-pAMPK (Thr-172) antibody, and the total SnRK1 protein was detected using the α-FLAG antibody. Total NPR1 protein was stained using Coomassie brilliant blue (CBB). *Non-specific band. (B) LC-MS/MS analysis of *in vitro* phosphorylated NPR1 by SnRK1. Peaks represent the intensity of the most abundant peptides containing pSer-557 detected in each group. (C) Interaction between GFP-fused SnRK1α and NPR1-mCherry or HA-mCherry transiently expressed in *N. benthamiana* after treatment with 1 mM SA for 6 h. (D) Interaction between endogenous SnRK1α and NPR1-GFP or GFP in transgenic *Arabidopsis* treated with water (-) or 0.3 mM SA (+) for 6 h. (E) *In vitro* interaction between SnRK1α or SnAK1 and GST-fused NPR1. (F) Effect of *snrk1* mutations on Ser-557 phosphorylation. NPR1-GFP was transiently expressed in WT and *snrk1α1^−/−^ snrk1α2^+/−^* (*sesquiα2-1* and *sesquiα2-2*) leaf-derived protoplasts, followed by treatment with mock or 0.3 mM SA for 6 h. NPR1-GFP was immunoprecipitated using the GFP-trap, and the enriched proteins were analyzed by western blotting with the α-pS557 and α-GFP antibodies. Numbers represent fold changes in Ser-557 phosphorylation induced by SA compared to the mock sample normalized to the total NPR1-GFP. (G) Effect of *snrk1* mutations on Ser-557 phosphorylation of the endogenous NPR1. Mature WT and *sesquiα2-1* plants were treated with mock (-) or 1 mM SA (+) for 8 h. NPR1 protein was enriched through immunoprecipitation using the α-NPR1 antibody and subsequently analyzed by western blotting using the α-pS557 antibody. Total NPR1 protein levels were measured with the α-NPR1 antibody. See also Figure S2.

**Figure S2.**
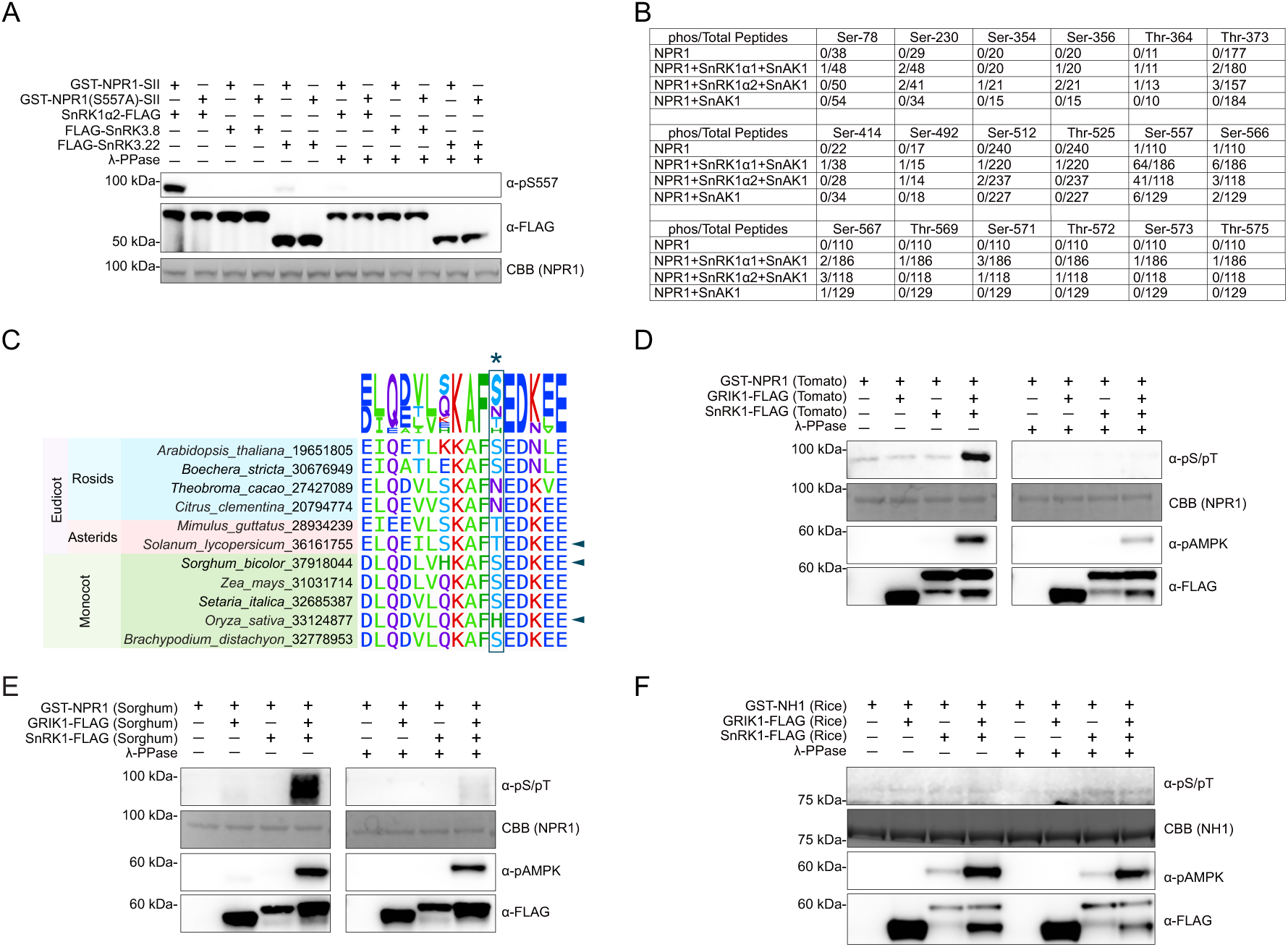
Phosphorylation of NPR1 by SnRK1 is largely conserved in plant species, related to Figure 3. (A) *In vitro* phosphorylation assays of recombinant NPR1. FLAG-tagged kinases were purified from *N. benthamiana*. Ser-557 phosphorylation was detected using the α-pS557 antibody. Total NPR1 protein was stained using CBB. (B) LC-MS/MS analysis of NPR1 phosphorylated *in vitro* by SnRK1. Related to Figure 3B. (C) Phylogenetic analysis of NPR1 from various plant species around the Ser-557 residue (marked by *). Sequence names in the figure are displayed as *Species name*_PACID. (D-F) *In vitro* phosphorylation assays of NPR1 by SnRK1 using cloned NPR1,GRIK1, and SnRK1 from Tomato (D), Sorghum (E), and Rice (F). NPR1 ortholog phosphorylation was detected using the α-pS/pT antibody, and SnRK1 activity was assessed using the α-pAMPK (Thr-172) antibody.

To investigate whether phosphorylation of NPR1 by SnRK1 is conserved across plant species, a phylogenetic study was performed, which showed that a serine or threonine residue corresponding to Ser-557 in AtNPR1 is largely conserved (Figure S2C). We found that in tomato (*Solanum lycopersicum*) and sorghum (*Sorghum bicolor*) orthologs, where the residues are conserved, NPR1 phosphorylation occurred with the application of SnRK1 activated by the AtSnAK ortholog GEMINIVIRUS REP-INTERACTING KINASE (GRIK) (Figures S2D and S2E). In contrast, OsNH1, the rice (*Oryza sativa*) ortholog with a histidine in the place of serine or threonine, was not phosphorylated by OsSnRK1 (Figure S2F).

### *SnRK1* is required for SA- and NPR1-mediated immunity

We then examined the induction of the immune-related genes in WT plants and in *sesquiα2* mutants using qPCR and found that both *sesquiα2* mutants were defective in the induction of SAR marker genes, *PR1* and *WRKY38* (Figures 4A and 4B). To comprehensively investigate the role of SnRK1 in SA-mediated immunity and its epistatic relationship with NPR1, we performed QuantSeq analysis on leaf samples of WT, *npr1-2*, *npr1^S557A^*, and both *sesquiα2* mutants, 24 hours after mock or SA treatment. Under mock conditions, all mutants exhibited transcriptional patterns closely resembling those of the WT, consistent with previous reports that SnRK1 retains partial functionality in the *sesquiα2* mutants^59,60^, and NPR1 is quiescent in the absence of the stimulus^17^ (Figures 4C and 4D). Upon SA treatment, the transcriptional landscape changed markedly with all mutants displaying patterns distinct from WT. Analysis revealed 3,666 SA-responsive differentially expressed genes in WT, compared to 1,865 in *npr1-2*, 2,241 in *npr1^S557A^*, and only 845 and 645 in *sesquiα2-1* and *sesquiα2-2*, respectively (|log2foldchange| > 1, adjusted p-value < 0.05) (Figures 4E and 4F). Gene Ontology (GO) term Biological Process (BP) and Kyoto Encyclopedia of Genes and Genomes (KEGG) analyses of SA-induced, Ser-557- and SnRK1-dependent genes showed enrichments for defense responses and protein secretory pathways (Figures 4G and 4H), processes previously demonstrated to be directly controlled by NPR1 and essential for SAR establishment^10^. Conversely, SA repressed, and Ser-557- and SnRK1-dependent genes were predominantly associated with plant growth (Figure 4I). Importantly, similar to the *npr1^S557A^* mutant (Figure 1F), the *sesquiα2* mutants were compromised in basal resistance as well as SA-induced defense against *Psm* ES4326 (Figure 4J).

**Figure 4.**
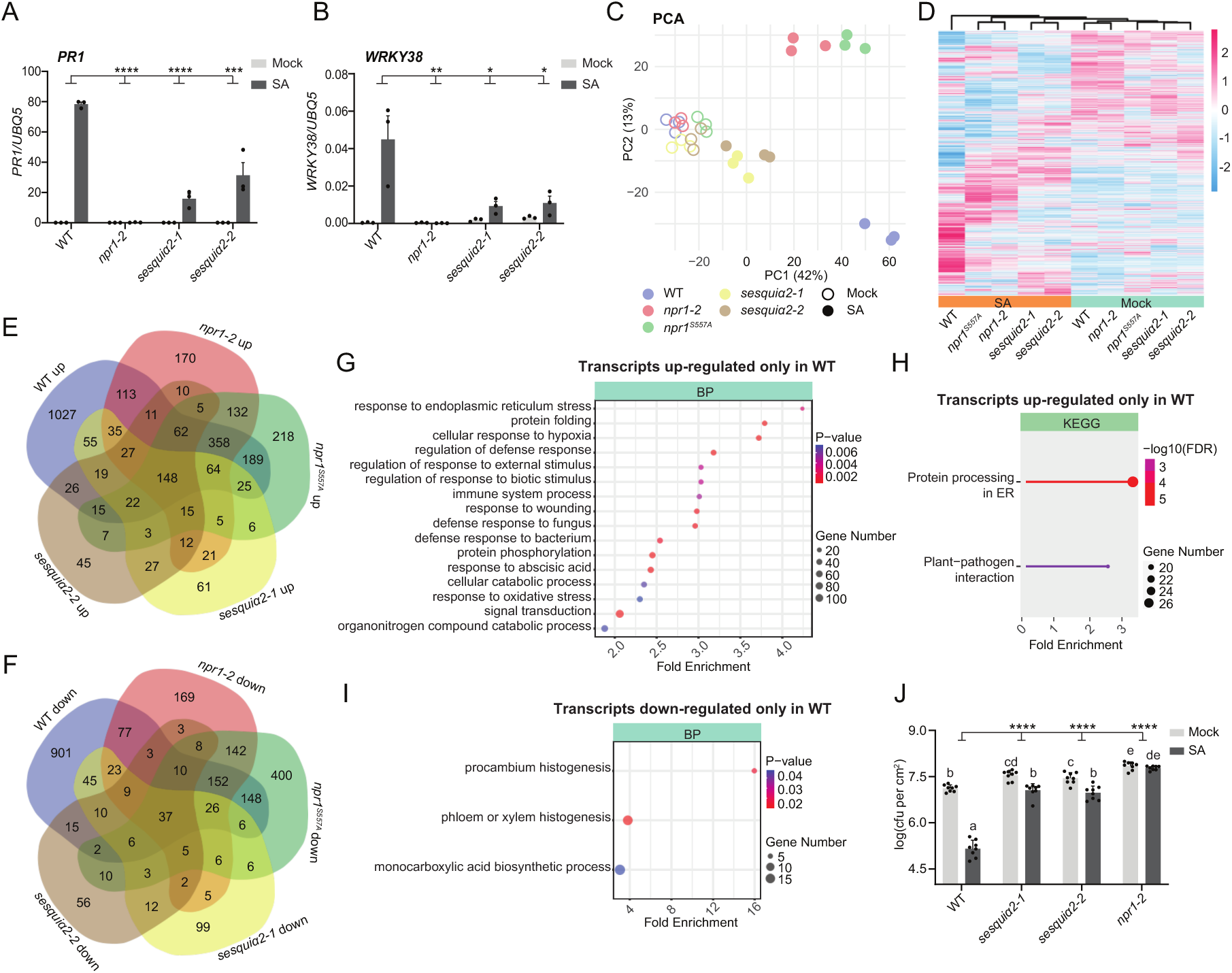
SnRK1 mediates SA-triggered immunity through NPR1. (A and B) *PR1* (A) and *WRKY38* (B) expression levels in 3.5-week-old WT, *npr1-2*, *sesquiα2-1* and *sesquiα2-2* plants 24 h after water or 1 mM SA treatment using RT-qPCR. (C and D) Principal component analysis (PCA) (C) and hierarchical clustering (D) of gene expression from QuantSeq analysis of 3.5-week-old WT, *npr1-2*, *sesquiα2-1*, and *sesquiα2-2* plants 24 h after treatment with water or 1 mM SA. (E and F) Venn diagrams comparing up-regulated (E) and down-regulated (F) differentially expressed genes across different genotypes. (G and H) Gene ontology (GO) term analysis of biological process (BP) (G) and Kyoto Encyclopedia of Genes and Genomes (KEGG) pathway analysis (H) of genes up-regulated only in WT. (I) GO term analysis of BP for genes down-regulated only in WT. (J) Basal and SA-induced resistance in WT, *sesquiα2-1*, *sesquiα2-2*, and *npr1-2* plants treated with mock or 1 mM SA 24 h before inoculation with *Psm* ES4326 (OD_600_ = 0.001). Bacterial growth was determined at 3 dpi. *n* = 8. All data are presented as mean ± s.e.m. Individual columns were compared using one-way ANOVA with Tukey’s post-hoc (J) and the interactions were tested using two-way ANOVA (A, B, and J). *p < 0.05, **p < 0.01, ***p < 0.001, ****p < 0.0001. n.s., not significant; different lowercase letters indicate statistical significance tested between multiple groups by one-way ANOVA at p < 0.05.

### SA coordinates NPR1-dependent immunity and NPR1-independent metabolism through activation of SnRK1

To investigate whether SA-induced phosphorylation of Ser-557 requires activation of SnRK1 in addition to triggering NPR1 nuclear translocation, we treated *NPR1-GFP* plants with the membrane-permeable reducing agent glutathione monoethyl ester (GSHmee) to directly reduce NPR1 from its oligomer, and also examined the transgenic lines expressing *npr1^C82A-^GFP* and *npr1^C216A^-GFP* with constitutive nuclear localization^17^. We found that nuclear localization of NPR1 alone was insufficient to induce Ser-557 phosphorylation (Figures S3A and S3B), suggesting the existence of distinct SA- and SnRK1-dependent mechanism to facilitate NPR1 phosphorylation.

**Figure S3.**
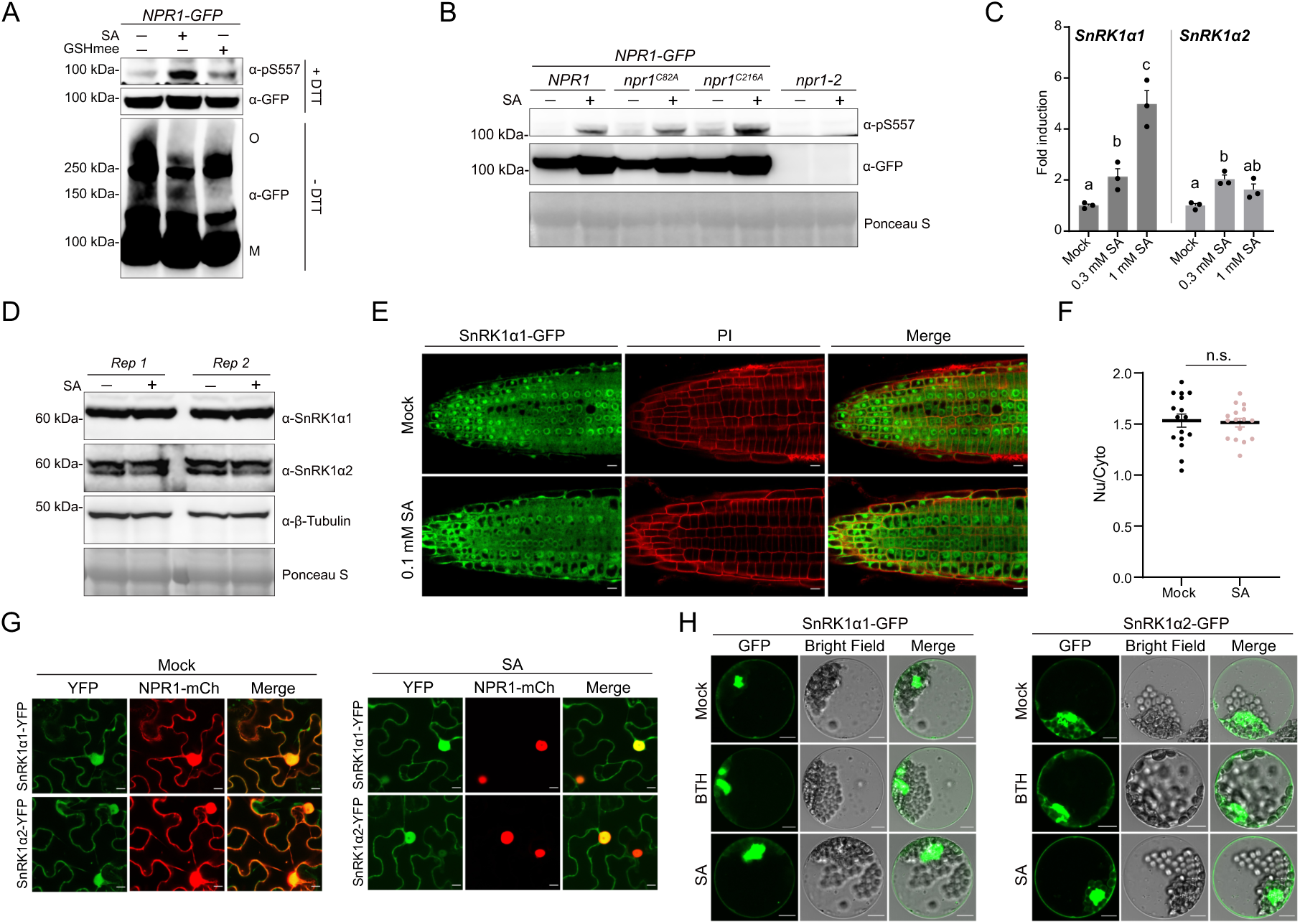
SA activates Ser-557 phosphorylation without affecting SnRK1 expression or subcellular localization, related to. **Figure 5**. (A) Effect of NPR1 nuclear translocation on Ser-557 phosphorylation. *35S:NPR1-GFP* seedlings were treated with mock, 0.3 mM SA, or 3 mM GSHmee for 6 h, then subjected to reducing (+50 mM DTT) or non-reducing (-DTT) SDS-PAGE, followed by western blot analysis. M, NPR1 mononer; O, NPR1 oligomer. (B) Analysis of the Ser-557 phosphorylation status of nuclear-localized NPR1. *35S:NPR1-GFP*, *35S:npr1^C82A^-GFP*, *35S:npr1^C216A^-GFP*, and *npr1-2* seedlings were treated with mock or 0.3 mM SA for 6 h, before analysis by western blotting. Equal loading was confirmed using Ponceau S staining. (C) *SnRK1α1* and *SnRK1α2* expression upon SA treatment measured through RT-qPCR. WT seedlings were treated with mock, 0.3 mM SA, or 1 mM SA for 6 h. *UBQ5* expression was used for normalization. (D) SnRK1α1 and SnRK1α2 protein levels upon SA treatment. WT seedlings were treated as in (C). Proteins were analyzed by western blotting using α-SnRK1α1, α-SnRK1α2, and α-β-tubulin antibodies. β-Tubulin was used as a loading control. Rep, biological replicate. (E and F) SnRK1α1 subcellular localization. Confocal micrographs of *35S:SnRK1α1-GFP* seedlings treated with mock or 0.1 mM SA for 6 h (E). Propidium iodide (PI) staining was used to visualize root outlines. Fluorescence intensity in (E) was measured using Fiji and quantified (F). Scale bars, 10 μm. (G) SnRK1α subcellular localization. Confocal micrographs of *N. benthamiana* transiently expressing YFP-tagged SnRK1α1 or SnRK1α2 and mCherry-tagged NPR1 at 30 h post-Agrobacterium infiltration. Mock or 1 mM SA treatments were applied 24 h post-infiltration. Scale bars, 10 μm. (H) SnRK1α subcellular localization. Confocal micrographs of *Arabidopsis* leaf-derived protoplasts transiently expressing GFP-tagged SnRK1α1 or SnRK1α2 and treated with mock, 0.1 mM Benzothiadiazole (BTH), or 0.3 mM SA for 6 h. Scale bars, 10 μm. All data are presented as mean ± s.e.m. Individual columns were compared using one-way ANOVA with Tukey’s post-hoc (C) or a two-tailed Student’s t-test (F). n.s., not significant; different lowercase letters indicate statistical significance tested between multiple groups by one-way ANOVA at p < 0.05.

Since SA treatment caused only a moderate increase in *SnRK1α1* and *SnRK1α2* transcripts, had a marginal effect on protein levels (Figures S3C and S3D), and did not alter subcellular localization (Figures S3E-S3H), we hypothesized that SA might impact Ser-557 phosphorylation by enhancing SnRK1 activity. Indeed, when we compared SA-induced genes from our QuantSeq experiments with the genes upregulated in response to transient *SnRK1α1* overexpression identified in a previous study^32^, we found a significant overlap (34.9% of the transcripts activated by *SnRK1α1* overexpression were also induced by SA), enriched in sugar starvation response and amino acid catabolism, suggesting that SA might activate SnRK1 (Figures S4A-S4C). To directly assess the effect of SA on SnRK1 activity *in vivo*, we measured nuclear phosphorylation of an SnRK1α-activity reporter, NLS-ratACC (Acetyl-CoA Carboxylase 1 peptide)-GFP-HA harboring the SnRK1/AMPK phosphorylation target sequence^61^. We observed a notable SA-induced increase in the WT SnRK1 background, but not in the *snrk1a1* mutant background (Figure 5A). Furthermore, SnRK1 target genes, *DARK INDUCIBLE 2* (*DIN2*), *DIN6*, and *SENESCENCE 5* (*SEN5*), which are upregulated under energy stress (e.g., low sucrose, low glucose, or extended darkness)^32,37^, were also strongly induced by SA in a dose-dependent manner (Figure 5B). Interestingly, this effect was not reliant on NPR1, as transcripts of *DIN6* and *SEN5* were still upregulated by SA in the *npr1-2* mutant in both seedlings (Figure S4D) and mature *Arabidopsis* plants (Figures S4E and S4F).

**Figure 5.**
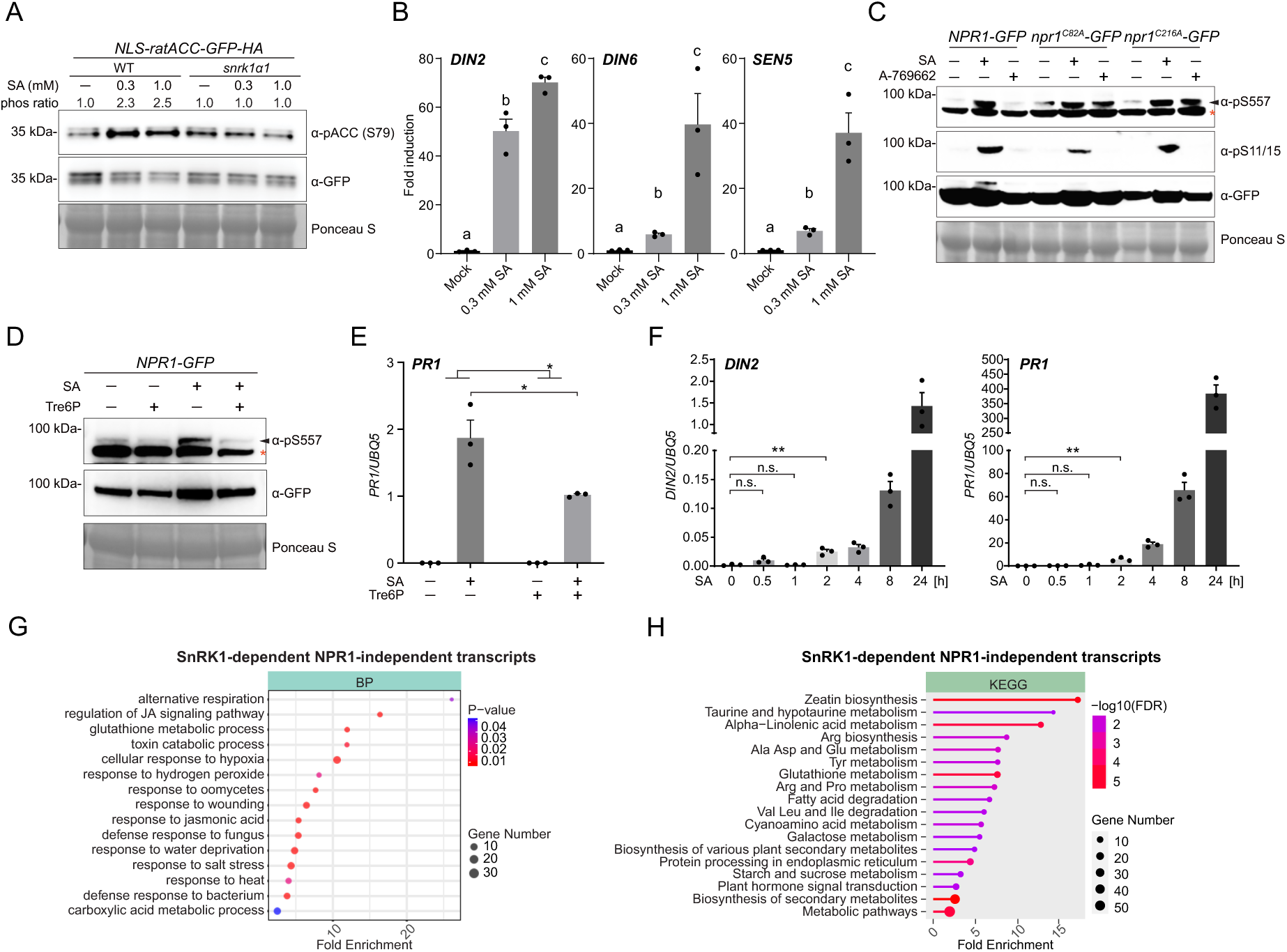
SA initiates defense and metabolic reprograming by modulating SnRK1 activity. (A) Effects of SA on SnRK1 activity. WT or *snrk1α1* seedlings carrying the *35S:NLS-ratACC-GFP-HA* reporter of SnRK1 activity were treated with mock, 0.3 mM SA, or 1 mM SA for 24 h. Nuclear SnRK1-dependent phosphorylation of the reporter was detected using the α-phospho-ACC (Ser-79) antibody, and total reporter protein levels were analyzed using the α-GFP antibody. Equal loading was confirmed using Ponceau S staining. (B) SnRK1 target genes *DIN2*, *DIN6*, and *SEN5* expression in response to SA measured by RT-qPCR. WT seedlings were treated with mock, 0.3 mM SA, or 1 mM SA for 24 h. *UBQ5* expression was used for normalization. (C) Effects of SnRK1 activation on NPR1 PTMs in the nucleus. *35S:NPR1-GFP*, *35S:npr1^C82A^-GFP* (in *npr1-2*), and *35S:npr1^C216A^-GFP* (in *npr1-2*) seedlings were treated with mock, 0.3 mM SA or 100 μM A-769662 for 6 h. Proteins were analyzed through western blotting using α-pS557, α-pS11/15, and α-GFP antibodies. *, nonspecific band. (D and E) Effects of SnRK1 inhibitor on Ser-557 phosphorylation and *PR1* expression. *35S:NPR1-GFP* seedlings grown in sucrose-free ½ MS medium were pretreated with or without 1 mM Trehalose 6-phosphate (Tre6P) for 2 h and subsequently treated with or without 0.3 mM SA for 6 h. Proteins were analyzed by western blotting using α-pS557 and α-GFP antibodies (D) and *PR1* expression levels were determined by RT-qPCR (E). *, nonspecific band. (F) Time course analysis of *PR1* and *DIN2* gene expression in *35S:NPR1-GFP* seedlings treated as in Figure 2B by RT-qPCR. *UBQ5* expression was used for normalization. (G and H) GO-term BP (G) and KEGG analyses (H) of 358 transcripts upregulated in WT, *npr1^S557A^-GFP*, and *npr1-2*, but not in *sesquiα2-1* or *sesquiα2-2*, upon SA treatment (Related to Figures 4D and 4E). All data are presented as mean ± s.e.m. Individual columns were compared using one-way ANOVA with Tukey’s post-hoc (B) or a two-tailed Student’s t-test (E and F), and the interactions were tested using two-way ANOVA (E). *p < 0.05, **p < 0.01. n.s., not significant; different lowercase letters indicate statistical significance tested between multiple groups by one-way ANOVA at p < 0.05. See also Figures S3 and S4.

**Figure S4.**
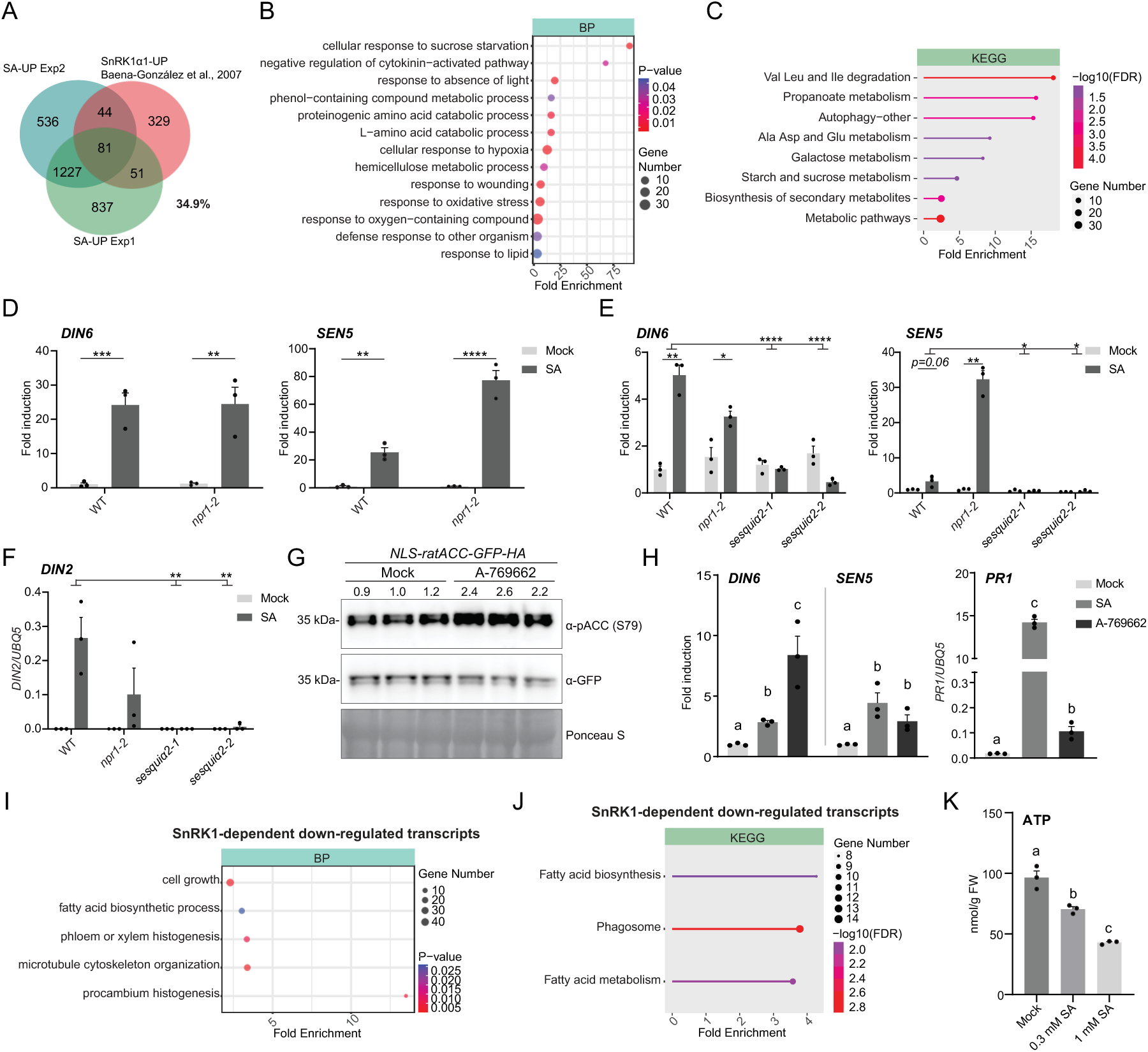
SnRK1 links SA-mediated defense and metabolic reprogramming, related to. **Figure 5**. (A-C) Venn diagram (A) comparing two independent QuantSeq datasets of SA-induced genes with a published transcriptional profile of genes up-regulated by SnRK1α1 activation^32^. GO-term BP (B) and KEGG pathway (C) analyses of 176 overlapping genes between the SA-induced and SnRK1α1 activated datasets. (D) Induction of SnRK1 target genes *DIN6* and *SEN5* expression in WT and *npr1-2* seedlings treated with mock or 1 mM SA for 24 h measured by RT-qPCR. *UBQ5* expression was used for normalization. (E) Induction of SnRK1 target genes expression in mature plants in response to SA, measured by RT-qPCR. *DIN6* and *SEN5* expression relative to mock in WT, *npr1-2, sesquiα2-1,* and *sesquiα2-2* 24 h after treatment with mock or 1 mM SA. *UBQ5* expression was used for normalization. (F) Induction of SnRK1 target gene *DIN2* expression in mature plants in response to SA, measured by RT-qPCR. Plants were treated as in (E). *UBQ5* expression was used for normalization. (G) Effects of A-769662 on the SnRK1-activity reporter. Seedlings carrying the *35S:NLS-ratACC-GFP-HA* reporter were treated with mock or 100 μM A-769662 for 8 h. The SnRK1 activity was measured using the α-phospho-ACC (Ser-79) antibody and the total reporter protein was analyzed using an α-GFP antibody. Three biological replicates are shown. Equal loading was confirmed using Ponceau S staining. (H) Expression of SnRK1 target genes *DIN6* and *SEN5* and NPR1 target gene *PR1* in response to A-769662 measured by RT-qPCR. WT seedlings were treated with mock or 100 μM A-769662 for 8 h. *UBQ5* expression was used for normalization. (I and J) GO-term BP (I) and KEGG (J) analyses of 152 transcripts downregulated in WT, *npr1^S557A^-GFP*, and *npr1-2*, but not in *sesquiα2-1* or *sesquiα2-2*, upon SA treatment. (K) ATP levels in WT seedlings after mock or SA treatment for 2 h. *n* = 3. All data are presented as mean ± s.e.m. Individual columns were compared using one-way ANOVA with Tukey’s post-hoc (H and K) or a two-tailed Student’s t-test (D and E), and the interactions were tested using two-way ANOVA (E and F). *p < 0.05,**p < 0.01,***p < 0.001,****p < 0.0001. Different lowercase letters indicate statistical significance tested between multiple groups by one-way ANOVA at p < 0.05.

We then tested whether elevated SnRK1 activity promotes Ser-557 phosphorylation using the AMPK activator A-769662^62^ which has been previously demonstrated to induce SnRK1 activity in rice^63^. As expected, A-769662 increased the nuclear SnRK1 activity, indicated by the increased phosphorylation of the NLS-ratACC reporter (Figure S4G). However, while A-769662 activated canonical SnRK1 outputs *DIN6* and *SEN5*, its effect on *PR1* transcript level was much lower compared to that in response to SA treatment, likely due to NPR1 being sequestered in the cytoplasm without redox changes triggered by SA^17,64^ (Figure S4H). Consistent with our hypothesis that SnRK1-mediated phosphorylation of Ser-557 is necessary but only partially sufficient for NPR1 activation, A-769662 treatment of *npr1^C82A^* and *npr1^C216A^* mutants with constitutive nuclear accumulation induced phosphorylation of Ser-557 but not Ser-11/15 (Figure 5C). Complementing these results, we found that treating plants with the SnRK1 inhibitor trehalose-6-phosphate (Tre6P) compromised SA-induced Ser-557 phosphorylation (Figure 5D) and dampened the *PR1* induction (Figure 5E). Collectively, these data indicate that Ser-557 phosphorylation requires not only SA-mediated reduction and nuclear translocation of NPR1, but also SA-induced activation of SnRK1.

We then investigated the timeline of NPR1 activation by SnRK1, using *DIN2* as a readout of SnRK1 activation, and *PR1* as an output of NPR1 induction. We found a strong correlation among SA-induced SnRK1 activation, Ser-557 phosphorylation, and NPR1 transcriptional activity, all of which showed significant induction starting at 2 hours after SA treatment (Figures 2B and 5F).

To better dissect the SA signaling independent of NPR1, we performed GO term and KEGG analyses on genes that are induced by SA in a SnRK1-dependent but NPR1-independent manner. Redox-and abiotic stress-related biological processes were enriched with the alternative respiration as the top GO term, while amino acid metabolism was enriched in the KEGG analysis (Figures 5G and 5H). In contrast, the SnRK1-dependent down-regulated processes included growth, metabolism, and cytoskeleton organization, while the KEGG analysis detected enrichment of fatty acid metabolism (Figures S4I and S4J). Finally, we found that treating plants with SA lowered the intracellular ATP levels in a dose-dependent manner (Figure S4K), suggesting a strong effect on the cellular energy state. Thus, although the direct molecular mechanism by which SA activates SnRK1 remains to be elucidated, our findings suggest that, through SnRK1, SA plays a dual role in orchestrating immune response and modulating the plant primary metabolic activities.

### SA inhibits TOR-mediated phosphorylation of NPR1 at Ser-55/59 in a SnRK1-dependent manner

The enrichment of multiple amino acid metabolic processes in our KEGG analysis of SA-induced SnRK1-dependent upregulated transcripts (Figure 5H) implies potential shifts in cellular translational activity. To evaluate the protein synthesis rate after SA treatment, we performed the surface sensing of translation (SUnSET) assay^65,66^, in which an analog of tyrosyl-tRNA, puromycin, is incorporated into the nascent peptides and measured by immunoblotting. We observed that SA treatment reduced the global translation efficiency to ∼50% within 4 hours in a SnRK1-dependent, but NPR1-independent, manner (Figure 6A).

**Figure 6.**
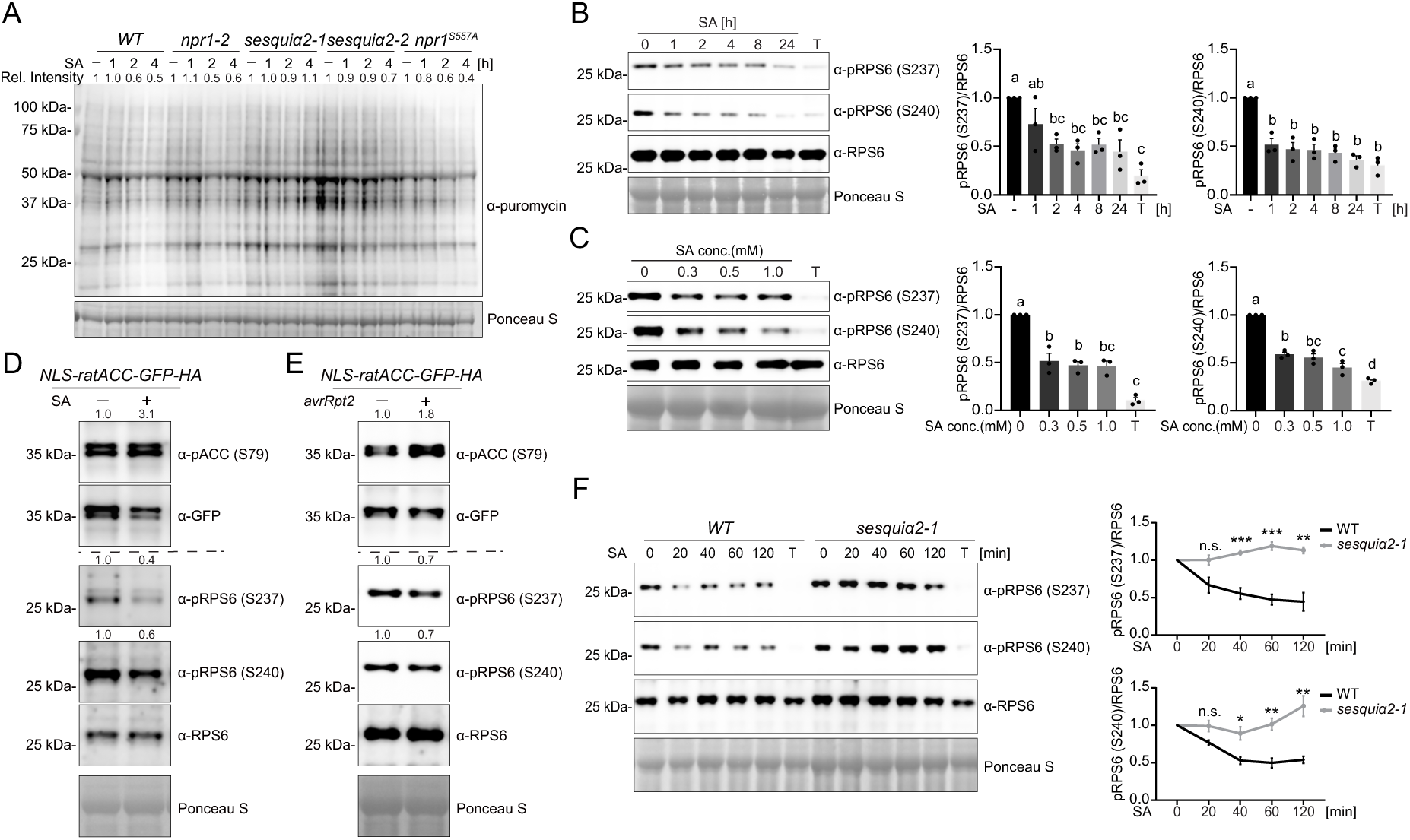
SA inhibits TOR activity through SnRK1. (A) SUnSET assay of nascent protein synthesis in response to SA in WT, *npr1^S557A^-GFP* (*npr1^S557A^*), *npr1-2, sesquiα2-1*, and *sesquiα2-2* plants. Four 5-mm leaf disks from 3.5-week-old plants were floated in water or 1 mM SA for indicated time periods. Puromycin labeling was initiated 45 min before sample collection and detected by western blotting using the α-puromycin antibody. Equal loading was confirmed using Ponceau S staining. (B) Time course of TOR-mediated RPS6 phosphorylation following SA treatment. WT seedlings were treated with 0.3 mM SA for the indicated durations or 10 μM Torin 2 (T) for 2 h. Proteins were western blotted (left) using the α-RPS6, α-pRPS6 (Ser-237), and α-pRPS6 (Ser-240) antibodies and band intensity was quantified (right) by normalizing to mock-treated samples. *n =* 3. (C) Effects of varying concentrations of SA on RPS6 phosphorylation. WT seedlings were treated with mock, 0.3, 0.5, and 1 mM SA, or 10 μM Torin 2 (T) for 2 h. Protein analysis and quantification were performed as in (B). (D) Effects of SA on SnRK1 activity and TOR-mediated RPS6 phosphorylation. Mature WT plants carrying *35S:NLS-ratACC-GFP-HA* were treated with mock (-) or 1 mM SA (+) for 24 h, then subjected to western blotting with α-pACC (Ser-79), α-RPS6, α-pRPS6 (Ser-237), α-pRPS6 (Ser-240), and α-GFP antibodies. (E) Effects of *Psm* ES4326/avrRpt2 (avrRpt2) on SnRK1 activity and TOR-mediated RPS6 phosphorylation. Mature WT plants carrying *35S:NLS-ratACC-GFP-HA* were infiltrated with mock (-) or *Psm* 4326/avrRpt2 (OD_600_ = 0.01) (+) in two local leaves. Samples from two systemic leaves were collected at 2 dpi. Protein analysis was performed as described in (D). (F) The role of SnRK1 in inhibiting TOR-mediated RPS6 phosphorylation during SA induction. WT and *sesquiα2-1* seedlings were treated with 0.3 mM SA for the indicated time or 10 μM Torin 2 (T) for 2 h. Western blotting (left) was performed as described in (B) and relative pPRS6 band intensities compared to mock-treated samples were plotted (right). *n* = 3. All data are presented as mean ± s.e.m. Individual columns were compared using one-way ANOVA with Tukey’s post-hoc (B and C) or a two-tailed Student’s t-test (F), *p < 0.05, **p < 0.01,***p < 0.001. n.s., not significant; different lowercase letters indicate statistical significance tested between multiple groups by one-way ANOVA at p < 0.05. See also Figure S5.

This result led us to investigate TOR, another central nutrient-sensing kinase, which not only acts antagonistically to SnRK1, but also regulates translation efficiency by phosphorylating its downstream substrate, P70-S6 Kinase (S6K)^67,68^. Both S6K and its direct target, ribosomal protein S6 (RPS6), are key components of cap-dependent translation, and their phosphorylation status is an established readout of TOR activity^30^. Therefore, we first assessed the phosphorylation of S6K and RPS6 in seedlings following SA treatment and found that SA rapidly inhibited the phosphorylation of both S6K (Figures S5A and S5B) and RPS6 (Figures 6B and 6C). SA also inhibited S6K phosphorylation in *Arabidopsis* leaf protoplasts (Figure S5C). Furthermore, the increase in SnRK1 activity was accompanied by inhibition of TOR that occurred not only in response to SA treatment in mature plants, but also in the distal systemic tissue after local ETI response triggered by *Psm* ES4326/*avrRpt2* (Figures 6D and 6E).

**Figure S5.**
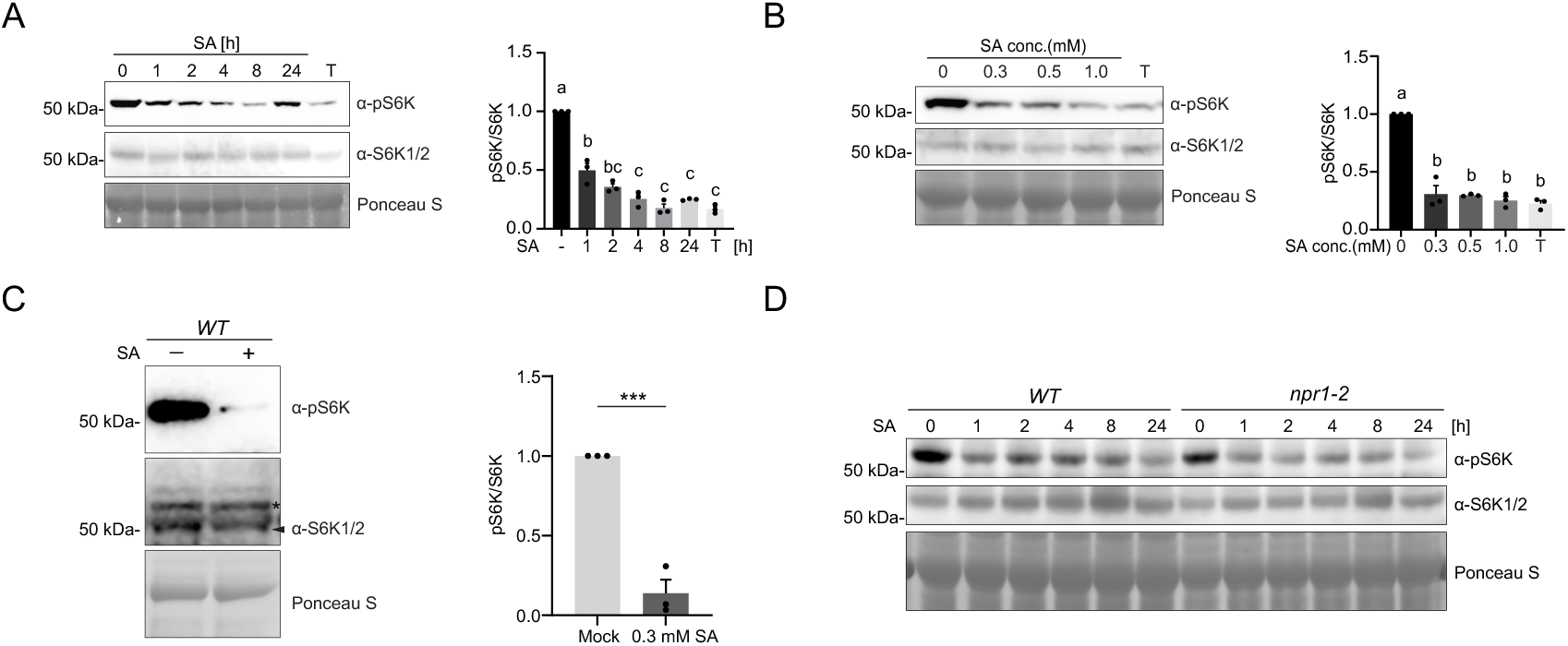
SA inhibits TOR activity, independent of NPR1, related to Figure 6. (A) Time course of TOR-mediated S6K phosphorylation following SA treatment. WT seedlings were treated with 0.3 mM SA for the indicated durations or 10 μM Torin 2 (T) for 2 h. Proteins were western blotted (left) using α-S6K1/2 and α-phospho-S6K (Thr-449) antibodies and band intensity was quantified (right) by normalizing to mock-treated samples. Equal loading was confirmed using Ponceau S staining. *n =* 3. (B) Effects of varying concentrations of SA on TOR-mediated S6K phosphorylation. WT seedlings were treated with varying concentrations of SA or 10 μM Torin 2 (T) for 2 h. Protein analysis and quantification were performed as in (A). (C) Effects of SA on S6K phosphorylation. WT *Arabidopsis* leaf-derived protoplasts were treated with mock or 0.3 mM SA for 2 h. Protein blotting (left) was performed as described in (A), and quantification (right). *Non-specific band (D) Roles of NPR1 in TOR-mediated S6K phosphorylation during SA induction. WT and *npr1-2* seedlings were treated with 0.3 mM SA for the indicated durations. Protein blotting was performed as described in (A). All data are presented as mean ± s.e.m. Individual columns were compared using one-way ANOVA with Tukey’s post-hoc (A and B) or a two-tailed Student’s t-test (C). ***p < 0.001. Different lowercase letters indicate statistical significance tested between multiple groups by one-way ANOVA at p < 0.05.

Since SA inhibits translation in a SnRK1 dependent manner, we examined whether TOR inhibition following SA treatment also depends on SnRK1. We compared RPS6 phosphorylation in WT and the *sesquiα2-1* mutant and observed that in *sesquiα2-1*, the SA-mediated decrease in RPS6 phosphorylation was absent over most of the sampling points in the 2-hour time-course (Figure 6F), suggesting that SnRK1 is required for the SA-mediated TOR inhibition. In contrast, the *npr1* mutant showed no effect on SA-triggered decrease in TOR phosphorylation of S6K, further confirming that SA-mediated inhibition of TOR activity occurs either upstream or independent of NPR1 (Figure S5D).

We next tested whether SA-mediated inhibition of TOR phosphorylation activity impacts NPR1 PTMs. We discovered that immunoprecipitated TOR complex could indeed phosphorylate NPR1 *in vitro*, and this effect was largely abolished by the TOR-specific inhibitor Torin 2 or by treatment with lambda phosphatase (Figure S6A). Since NPR1 is known to be phosphorylated at Ser-55/59 in the cytosol in its quiescent state, and this phosphorylation decreases shortly after exposure to SA, it’s possible that TOR might be the enzyme that targets Ser-55/59. Consistent with this hypothesis, the Ser-55/59 residues are highly conserved across angiosperms with the +1 residue of Ser-59 being a proline that matches the consensus sequence for TOR/mTOR substrates^69,70^ (Figure S6B). Using the phospho-specific antibody against Ser-55/59, we found that TOR indeed phosphorylated Ser-55/59, and this modification was inhibited by the addition of Torin 2 (Figure 7A). Moreover, the phosphorylation signal was markedly reduced in the Ser-55/59 mutant of NPR1 in the *in vitro* phosphorylation assay (Figure S6C). To validate TOR as a *bona fide* kinase of NPR1 *in vivo*, we treated *NPR1-GFP* transgenic seedlings and *NPR1-GFP*-transfected protoplasts with TOR-specific inhibitors Torin 2 or AZD-8055 and found that Ser-55/59 phosphorylation was abolished in both tests (Figures 7B and S6D). *In vivo* phosphorylation of Ser-55/59 by TOR was validated using an estradiol-inducible TOR silencing line (*tor-es*)^28,29^ (Figure 7C). Co-IP assay further validated that NPR1 interacts with the endogenous TOR, verifying the *in vivo* NPR1-TOR interaction (Figure 7D). To better understand the relationship between the activity of TOR, its interaction with NPR1, and Ser-55/59 phosphorylation, we immunoprecipitated NPR1-GFP and probed the sample with α-TOR antibody in a 2-hour time course after SA treatment. Simultaneously, we measured the RPS6 phosphorylation as an output of TOR activity as well as Ser-55/59 phosphorylation. We found that SA concurrently decreases TOR activity and abolishes its interaction with NPR1, possibly due to NPR1 nuclear translocation, as early as 30 minutes after SA treatment, explaining the rapid dephosphorylation of Ser-55/59 (Figure 7E).

**Figure 7.**
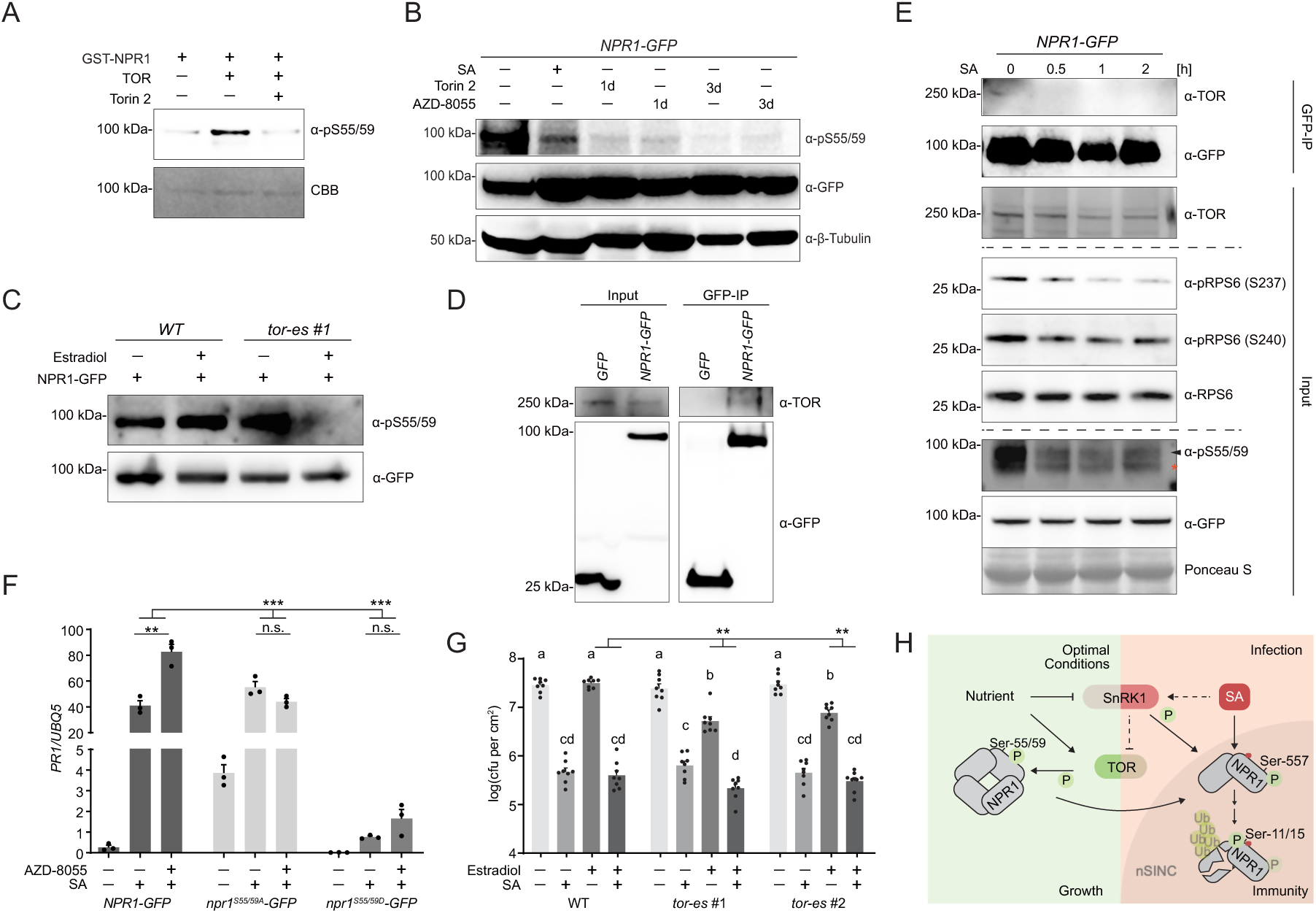
TOR phosphorylates NPR1 at Ser-55/59 under optimal conditions. (A) *In vitro* phosphorylation of recombinant NPR1 by TOR. Western blotting was performed using the α-pS55/59 antibody. Total NPR1 protein was stained using CBB. (B) Effect of TOR inhibition on Ser-55/59 phosphorylation. *35S:NPR1-GFP* seedlings were treated with mock, 0.3 mM SA (6 h), 10 μM Torin 2 (1 day or 3 days), or 2 μM AZD-8055 (1 day or 3 days). Phosphorylated NPR1 was detected using the α-pS55/59 antibody. Total NPR1 protein levels were determined using the α-GFP antibody. β-tubulin was used as a loading control. (C) Estradiol-induced silencing of *TOR* on Ser-55/59 phosphorylation. NPR1-GFP was transiently expressed in WT or *tor-es* leaf-derived protoplasts and treated overnight with either DMSO (mock) or 5 μM estradiol. Western blotting was performed as in (B). (D) Interaction of endogenous TOR with GFP-fused NPR1 or GFP in transgenic *Arabidopsis*. (E) Effects of SA on TOR activity and TOR-NPR1 interaction. *35S:NPR1-GFP* seedlings were treated with mock or 0.3 mM SA for the indicated durations. The lysate was split into two portions: one subjected to GFP immunoprecipitation followed by western blotting with α-TOR and α-GFP antibodies; and the other analyzed directly by western blotting as input using α-TOR, α-pRPS6 (Ser-237), α-pRPS6 (Ser-240), α-RPS6, α-pS55/59, and α-GFP antibodies. *Non-specific band. (F) Effect of TOR inhibition on *PR1* gene expression. RT-qPCR was performed on *35S:NPR1-GFP, 35S:npr1^S55/59A^-GFP*, and *35S:npr1^S55/59D^-GFP* seedlings pre-treated with mock or 2 μM AZD-8055 and subsequently treated with mock or 0.3 mM SA for 8 h. *UBQ5* expression was used for normalization. (G) Basal and SA-induced resistance in *tor* mutants. WT, *tor-es #1*, and *tor-es #2* plants were pre-treated with mock or 10 μM estradiol two days before mock or 1 mM SA application. One day after mock of SA treatment, *Psm* ES4326 (OD_600_ = 0.001) was inoculated and bacterial growth was determined at 3 dpi. *n* = 8. (H) Proposed model in which central metabolic regulators SnRK1 and TOR govern plant immunity through differential phosphorylation of NPR1, uncovering a mechanistic link to SA signaling and NPR1 activation in coordinating defense and growth. Under normal growth conditions (left), NPR1 is phosphorylated at Ser-55/59 by active TOR (bright green) and sequestered in the cytoplasm via oligomerization, while SnRK1 (dim red) is inhibited by nutrient availability. During immune responses (right), SA activates SnRK1 (bright red), leading to TOR inhibition and Ser-55/59 dephosphorylation. SA-triggered redox changes also facilitate nuclear translocation of NPR1. SnRK1 then phosphorylates NPR1 at Ser-557, partially activating it and priming it for subsequent PTMs. Ub, ubiquitination. All data are presented as mean ± s.e.m. Individual columns were compared using one-way ANOVA with Tukey’s post-hoc (G) or a two-tailed Student’s t-test (F), and the interactions were tested using two-way ANOVA (F and G). **p < 0.01,***p < 0.001. n.s., not significant; different lowercase letters indicate statistical significance tested between multiple groups by one-way ANOVA at p < 0.05. See also Figure S6.

**Figure S6.**
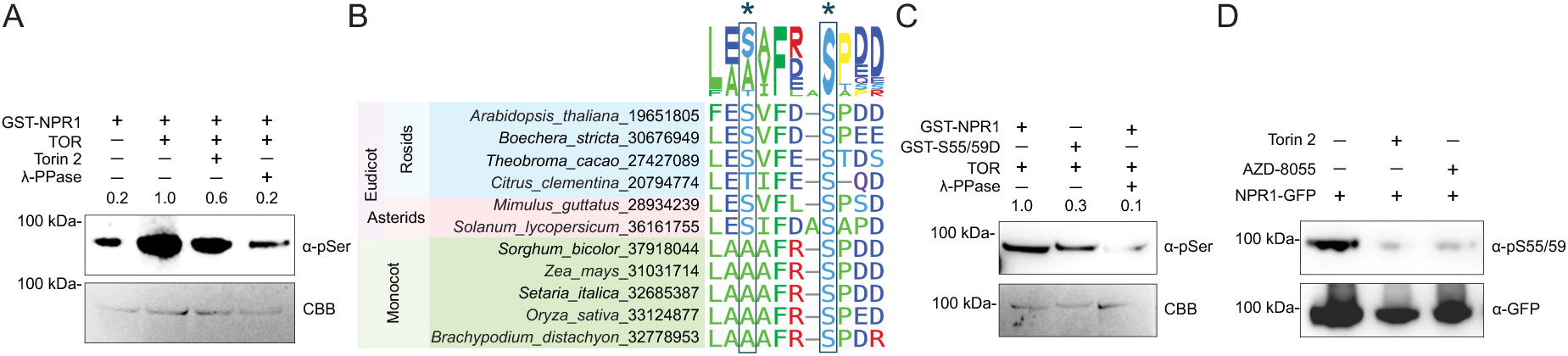
TOR phosphorylates NPR1 both *in vitro* and *in vivo*, related to Figure 7. (A) *In vitro* phosphorylation of recombinant NPR1 by immunoprecipitated TOR from *Arabidopsis* seedlings. Western blotting was performed using the α-pSer antibody. Total NPR1 protein was stained using CBB. (B) Phylogenetic analysis of NPR1 orthologs in various plant species around the Ser-55/59 residues (marked by *). Sequence names in the figure are displayed as *Species name*_PACID. (C) *In vitro* phosphorylation of recombinant NPR1 and *npr1^S55/59D^* by TOR. (D) *In vivo* phosphorylation of NPR1 at Ser-55/59. NPR1-GFP constructs were transiently expressed in WT *Arabidopsis* leaf-derived protoplasts and treated with DMSO (-), 10 μM Torin 2, or 2 μM AZD-8055 overnight. Western blotting was performed using the α-pS55/59 antibody.

To connect TOR activity with Ser-55/59-mediated immunity, we treated *NPR1-GFP* and its Ser-55/59 mutants with mock, SA, or a combination of SA and the TOR inhibitor AZD-8055. We found that inhibiting TOR with AZD-8055 further enhanced *PR1* induction by SA in the WT, and this effect was abolished in the Ser-55/59 phospho-mutants, supporting that TOR regulates *PR1* expression through modification of Ser-55/59.

Finally, to examine the functional impact of TOR on plant defense, we used estradiol, SA, and combined SA+estradiol treatments in two *tor-es* lines to assess the effect of Ser-55/59 dephosphorylation (Figure 7C) on both basal and SA-induced immunity. We found that silencing TOR enhanced basal immunity relative to the WT control (Figure 7G). However, SA treatment of TOR-silenced plants did not result in a greater defense response than that observed in WT, likely because knocking down *TOR* effectively mimics the SA-induced dephosphorylation of Ser-55/59 in NPR1.

## DISCUSSION

Despite the discovery of a number of NPR1 PTMs, with effects ranging from controlling NPR1 subcellular localization^16,22^ to its interaction with TFs^16^, the key modifications and enzymes involved in controlling NPR1’s function as a transcriptional coactivator remained unclear. In this study, we identified Ser-557 phosphorylation as a critical step in initiating and activating NPR1’s immune function. While Ser-557 phosphorylation does not affect its direct interactions with TGA3 TF and NIMIN1-3, the phospho-mimetic npr1^S557D^ mutant constitutively forms nSINCs, becomes transcriptionally active, and exhibits an auto-immune phenotype. In addition, Ser-557 phosphorylation occurs prior to the other known PTMs in the nucleus. Together, our data indicate that Ser-557 phosphorylation is a necessary and, likely, initial step in triggering transcriptional reprogramming and SAR. However, Ser-557 phosphorylation alone does not induce full-scale immunity, as *npr1^S557D^* remains sensitive to SA-induction. This is likely due to the absence of subsequent PTMs in this signaling cascade involving phosphorylation at Ser-11/15 by an unknown SA-dependent kinase and sequential ubiquitination.

The discovery of the two master nutrient regulators, SnRK1 and TOR, as pivotal kinases regulating NPR1 activity not only fills a critical gap in the signaling pathway between SA and NPR1 but also establishes a direct connection between SA-mediated immune responses and cellular metabolic processes essential for plant survival under biotic stress (Figure 7H). Phosphorylation of NPR1 at Ser-55/59 by TOR and oligomerization through intermolecular disulfide bond formation are dual mechanisms of keeping NPR1 as an inactive oligomer under normal conditions. Within an hour after SA treatment, SnRK1-dependent TOR inhibition occurs, and NPR1 interaction with TOR is disrupted, explaining the rapid decrease in phosphorylation at Ser-55/59. However, whether a phosphatase is actively involved in this process remains unknown. Upon translocation into the nucleus, NPR1 is phosphorylated at Ser-557 by SnRK1, partially activating it and priming it for additional PTMs. This shows that NPR1 activation is a multi-layered, stepwise process.

As part of their role in coordinating growth and stress responses, SnRK1 and TOR have been shown to antagonize each other under both biotic and abiotic stresses across species^38^. Interestingly, acetylsalicylate (aspirin), a derivative of salicylic acid, has been shown to activate AMPK in cancer cell lines while inhibiting mTOR signaling in a partially AMPK-dependent manner^71^. The antagonism between AMPK and mTOR is also present in the mammalian immune response against pathogens^72,73^. Differential phosphorylation of shared substrates by AMPK and mTOR has also been reported in mammals^74^. Under nutrient rich conditions, mTOR prevents UNC-51-LIKE KINASE 1 (ULK1) from initiating autophagy by phosphorylating Ser-757, while AMPK activation during glucose starvation facilitates autophagy through phosphorylation of Ser-317/777. This antagonistic regulation of autophagy is also conserved in plants, mediated by the direct phosphorylation of SnRK1 substrates^75,76^, a process that can be induced by SA and is required for restricting cell death during pathogen infection through NPR1^50^.

Identifying the mechanism by which SA activates SnRK1 is challenging, because SnRK1/AMPK responds broadly to diverse stresses, implying that these signals converge on a common upstream activator. In mammals, AMPK is triggered by an increased AMP-to-ATP ratio following cellular energy depletion^77^. However, plant SnRK1 orthologs are insensitive to AMP and ADP^78^, leaving their direct activation cue a mystery. Revisiting established triggers of SnRK1, such as extended darkness and hypoxia, reveals a unifying feature of energy deprivation^32^. Notably, SA reduces intracellular sugars^79^ and lowers intracellular ATP levels (Figure S4K) by inducing alternative respiration^23,80^, suggesting that SA-induced resource depletion might activate SnRK1.

As a key regulator of cellular stress responses, SnRK1/AMPK/SNF1 helps cells overcome adverse conditions through enzymatic regulation and transcriptional reprogramming^81,82^. The phosphorylation of NPR1 by SA-activated SnRK1 is an integral step in combating pathogen infection. Moreover, SA-induced, SnRK1-dependent genes are enriched in pathways related to the catabolism of sugars, amino acids, and fatty acids, emphasizing the importance of providing alternative energy sources and metabolites in response to biotic stress. Conversely, we show that SA inhibits the activity of the other master metabolic regulator, TOR, in a SnRK1-dependent, but NPR1-independent manner, leading to reduced cap-dependent translation as part of the growth-to-defense transition. Our results align with a recent study on SA repressing *Arabidopsis* growth under nutrient-rich conditions by interfering with the TOR pathway^83^, extending mechanistic insight into the previously reported translation inhibition in response to SA treatment^84,85,86^.

In summary, this study reveals three interconnected layers of SA regulation on plant physiology (Figure 7H). First, upon pathogen challenge, SA activates SnRK1, which inhibits TOR signaling to trigger the general growth-to-defense transition. Second, the rebalancing of activities between these two master kinases shifts NPR1 phosphorylation from Ser-55/59 to Ser-557. Third, this shift in phosphorylation leads to NPR1 activation and its subsequent PTMs to control the plant immune response in coordination with the cellular metabolic state.

## Limitations of the study

Our findings uncover a dynamic molecular network in which SA activates SnRK1 while inhibiting TOR. However, it remains unclear whether SA activates SnRK1 through a signal distinct from other stress response pathways or via a more general stress-induced activator. We found that the SA-mediated increases in SnRK1 transcriptional markers *DIN6* and *SEN5* were comparable to ABA-mediated induction, but significantly lower than the levels observed following extended darkness^37^. These results indicate that that SnRK1 activation exhibits variable degrees of induction depending on the type of stress. Furthermore, while SnRK1 is required for inhibiting TOR at early time points after SA treatment, the specific molecular mechanism remains unknown. Additionally, the full metabolic impact induced by SA in balancing SnRK1 and TOR has yet to be elucidated.

## Acknowledgements

We thank Dr. Elena Baena-Gonzalez for sharing the *sesquiα2* mutant lines; Dr. Csaba Koncz for the SnRK1α1-GFP line; Dr. Albrecht G. von Arnim and Abigail Burke for the α-RPS6, α-pRPS6 (Ser237), and α-pRPS6 (Ser240) antibodies. We thank Dr. Pawan Dahal for assisting with FPLC and protein purification; Duke University School of Medicine for the use of the Proteomics and Metabolomics Core Facility. We thank Dr. Jinlong Wang and Dong laboratory members for helpful discussions. This work was supported by grants from the National Institutes of Health (R35-GM118036-06); National Science Foundation (IOS-2041378), and the Howard Hughes Medical Institute to X.D; Deutsche Forschungsgemeinschaft (DFG; DR273/18-2) to W.D.L.

## Author Contributions

J.W., T.C., and X.D. conceived the project. Y.C., J.W., T.C., and S.K. designed the research. While J.W. performed the initial mass spectrometry to identify the NPR1 phospho-residue, generated phospho-mutant lines, tested their defense phenotypes, and conducted yeast two-hybrid assays, Y.C. carried out most of the subsequent experiments. T.C. assisted with the SUnSET assay and QuantSeq, and performed the bioinformatic analysis of the QuantSeq data. J.D. generated the NLS-ratACC-GFP-HA/*snrk1α1* reporter line. Y.X. performed the phylogenic analyses. W.D.L. supervised J.D. and provided feedback on the project and the manuscript. Y.C., S.K. and X.D. prepared the manuscript with the participation of all other authors.

## Materials availability

All unique constructs and reagents in this study will be available from the lead contact upon completion of Materials Transfer Agreement.

## References

1. Boller, T., and Felix, G. (2009). A renaissance of elicitors: perception of microbe-associated molecular patterns and danger signals by pattern-recognition receptors. Annu Rev Plant Biol 60, 379–406. 10.1146/annurev.arplant.57.032905.105346.

2. DeFalco, T.A., and Zipfel, C. (2021). Molecular mechanisms of early plant pattern-triggered immune signaling. Mol Cell 81, 4346. 10.1016/j.molcel.2021.09.028.

3. Cui, H., Tsuda, K., and Parker, J.E. (2015). Effector-triggered immunity: from pathogen perception to robust defense. Annu Rev Plant Biol 66, 487–511. 10.1146/annurev-arplant-050213-040012.

4. Jones, J.D., and Dangl, J.L. (2006). The plant immune system. Nature 444, 323–329. 10.1038/nature05286.

5. Malamy, J., Carr, J.P., Klessig, D.F., and Raskin, I. (1990). Salicylic Acid: a likely endogenous signal in the resistance response of tobacco to viral infection. Science 250, 1002–1004. 10.1126/science.250.4983.1002.

6. Cao, L., Karapetyan, S., Yoo, H., Chen, T., Mwimba, M., Zhang, X., and Dong, X. (2024). H(2)O(2) sulfenylates CHE, linking local infection to the establishment of systemic acquired resistance. Science 385, 1211–1217. 10.1126/science.adj7249.

7. Fu, Z.Q., and Dong, X. (2013). Systemic acquired resistance: turning local infection into global defense. Annu Rev Plant Biol 64, 839–863. 10.1146/annurev-arplant-042811-105606.

8. Cao, H., Bowling, S.A., Gordon, A.S., and Dong, X. (1994). Characterization of an Arabidopsis Mutant That Is Nonresponsive to Inducers of Systemic Acquired Resistance. Plant Cell 6, 1583–1592. 10.1105/tpc.6.11.1583.

9. Cao, H., Glazebrook, J., Clarke, J.D., Volko, S., and Dong, X. (1997). The Arabidopsis NPR1 gene that controls systemic acquired resistance encodes a novel protein containing ankyrin repeats. Cell 88, 57–63. 10.1016/s0092-8674(00)81858-9.

10. Wang, D., Weaver, N.D., Kesarwani, M., and Dong, X.N. (2005). Induction of protein secretory pathway is required for systemic acquired resistance. Science 308, 1036–1040. 10.1126/science.1108791.

11. Zhou, P., Zavaliev, R., Xiang, Y., and Dong, X. (2023). Seeing is believing: Understanding functions of NPR1 and its paralogs in plant immunity through cellular and structural analyses. Curr Opin Plant Biol 73, 102352. 10.1016/j.pbi.2023.102352.

12. Fan, W., and Dong, X. (2002). In vivo interaction between NPR1 and transcription factor TGA2 leads to salicylic acid-mediated gene activation in Arabidopsis. Plant Cell 14, 1377–1389. 10.1105/tpc.001628.

13. Zhang, Y., Fan, W., Kinkema, M., Li, X., and Dong, X. (1999). Interaction of NPR1 with basic leucine zipper protein transcription factors that bind sequences required for salicylic acid induction of the PR-1 gene. Proc Natl Acad Sci U S A 96, 6523–6528. 10.1073/pnas.96.11.6523.

14. Powers, J., Zhang, X., Reyes, A.V., Zavaliev, R., Ochakovski, R., Xu, S.L., and Dong, X. (2024). Next-generation mapping of the salicylic acid signaling hub and transcriptional cascade. Mol Plant 17, 1558–1572. 10.1016/j.molp.2024.08.008.

15. Withers, J., and Dong, X. (2016). Posttranslational Modifications of NPR1: A Single Protein Playing Multiple Roles in Plant Immunity and Physiology. PLoS Pathog 12, e1005707. 10.1371/journal.ppat.1005707.

16. Saleh, A., Withers, J., Mohan, R., Marques, J., Gu, Y., Yan, S., Zavaliev, R., Nomoto, M., Tada, Y., and Dong, X. (2015). Posttranslational Modifications of the Master Transcriptional Regulator NPR1 Enable Dynamic but Tight Control of Plant Immune Responses. Cell Host Microbe 18, 169–182. 10.1016/j.chom.2015.07.005.

17. Mou, Z., Fan, W., and Dong, X. (2003). Inducers of plant systemic acquired resistance regulate NPR1 function through redox changes. Cell 113, 935–944. 10.1016/s0092-8674(03)00429-x.

18. Tada, Y., Spoel, S.H., Pajerowska-Mukhtar, K., Mou, Z., Song, J., Wang, C., Zuo, J., and Dong, X. (2008). Plant immunity requires conformational changes [corrected] of NPR1 via S-nitrosylation and thioredoxins. Science 321, 952–956. 10.1126/science.1156970.

19. Spoel, S.H., Mou, Z., Tada, Y., Spivey, N.W., Genschik, P., and Dong, X. (2009). Proteasome-mediated turnover of the transcription coactivator NPR1 plays dual roles in regulating plant immunity. Cell 137, 860–872. 10.1016/j.cell.2009.03.038.

20. Skelly, M.J., Furniss, J.J., Grey, H., Wong, K.W., and Spoel, S.H. (2019). Dynamic ubiquitination determines transcriptional activity of the plant immune coactivator NPR1. Elife 8. 10.7554/eLife.47005.

21. Xie, C., Zhou, X., Deng, X., and Guo, Y. (2010). PKS5, a SNF1-related kinase, interacts with and phosphorylates NPR1, and modulates expression of WRKY38 and WRKY62. J Genet Genomics 37, 359–369. 10.1016/S1673-8527(09)60054-0.

22. Lee, H.J., Park, Y.J., Seo, P.J., Kim, J.H., Sim, H.J., Kim, S.G., and Park, C.M. (2015). Systemic Immunity Requires SnRK2.8-Mediated Nuclear Import of NPR1 in Arabidopsis. Plant Cell 27, 3425–3438. 10.1105/tpc.15.00371.

23. Rhoads, D.M., and McIntosh, L. (1992). Salicylic Acid Regulation of Respiration in Higher Plants: Alternative Oxidase Expression. Plant Cell 4, 1131–1139. 10.1105/tpc.4.9.1131.

24. Raskin, I., Turner, I.M., and Melander, W.R. (1989). Regulation of heat production in the inflorescences of an Arum lily by endogenous salicylic acid. Proc Natl Acad Sci U S A 86, 2214–2218. 10.1073/pnas.86.7.2214.

25. Vishwakarma, A., Tetali, S.D., Selinski, J., Scheibe, R., and Padmasree, K. (2015). Importance of the alternative oxidase (AOX) pathway in regulating cellular redox and ROS homeostasis to optimize photosynthesis during restriction of the cytochrome oxidase pathway in Arabidopsis thaliana. Ann Bot 116, 555–569. 10.1093/aob/mcv122.

26. Chen, Z., and Klessig, D.F. (1991). Identification of a soluble salicylic acid-binding protein that may function in signal transduction in the plant disease-resistance response. Proc Natl Acad Sci U S A 88, 8179–8183. 10.1073/pnas.88.18.8179.

27. Chen, Z., Silva, H., and Klessig, D.F. (1993). Active oxygen species in the induction of plant systemic acquired resistance by salicylic acid. Science 262, 1883–1886. 10.1126/science.8266079.

28. Xiong, Y., and Sheen, J. (2012). Rapamycin and glucose-target of rapamycin (TOR) protein signaling in plants. J Biol Chem 287, 2836–2842. 10.1074/jbc.M111.300749.

29. Xiong, Y., McCormack, M., Li, L., Hall, Q., Xiang, C., and Sheen, J. (2013). Glucose-TOR signalling reprograms the transcriptome and activates meristems. Nature 496, 181–186. 10.1038/nature12030.

30. Liu, Y., Hu, J., Duan, X., Ding, W., Xu, M., and Xiong, Y. (2025). Target of Rapamycin (TOR): A Master Regulator in Plant Growth, Development, and Stress Responses. Annu Rev Plant Biol. 10.1146/annurev-arplant-083123-050311.

31. Baena-Gonzalez, E., and Hanson, J. (2017). Shaping plant development through the SnRK1-TOR metabolic regulators. Curr Opin Plant Biol 35, 152–157. 10.1016/j.pbi.2016.12.004.

32. Baena-Gonzalez, E., Rolland, F., Thevelein, J.M., and Sheen, J. (2007). A central integrator of transcription networks in plant stress and energy signalling. Nature 448, 938–942. 10.1038/nature06069.

33. Jamsheer, K.M., Kumar, M., and Srivastava, V. (2021). SNF1-related protein kinase 1: the many-faced signaling hub regulating developmental plasticity in plants. J Exp Bot 72, 6042–6065. 10.1093/jxb/erab079.

34. Meng, Y., Zhang, N., Li, J., Shen, X., Sheen, J., and Xiong, Y. (2022). TOR kinase, a GPS in the complex nutrient and hormonal signaling networks to guide plant growth and development. J Exp Bot 73, 7041–7054. 10.1093/jxb/erac282.

35. Schepetilnikov, M., Dimitrova, M., Mancera-Martinez, E., Geldreich, A., Keller, M., and Ryabova, L.A. (2013). TOR and S6K1 promote translation reinitiation of uORF-containing mRNAs via phosphorylation of eIF3h. EMBO J 32, 1087–1102. 10.1038/emboj.2013.61.

36. Cho, H.Y., Wen, T.N., Wang, Y.T., and Shih, M.C. (2016). Quantitative phosphoproteomics of protein kinase SnRK1 regulated protein phosphorylation in Arabidopsis under submergence. J Exp Bot 67, 2745–2760. 10.1093/jxb/erw107.

37. Rodrigues, A., Adamo, M., Crozet, P., Margalha, L., Confraria, A., Martinho, C., Elias, A., Rabissi, A., Lumbreras, V., Gonzalez-Guzman, M., et al. (2013). ABI1 and PP2CA phosphatases are negative regulators of Snf1-related protein kinase1 signaling in Arabidopsis. Plant Cell 25, 3871–3884. 10.1105/tpc.113.114066.

38. Margalha, L., Confraria, A., and Baena-Gonzalez, E. (2019). SnRK1 and TOR: modulating growth-defense trade-offs in plant stress responses. J Exp Bot 70, 2261–2274. 10.1093/jxb/erz066.

39. Hulsmans, S., Rodriguez, M., De Coninck, B., and Rolland, F. (2016). The SnRK1 Energy Sensor in Plant Biotic Interactions. Trends Plant Sci 21, 648–661. 10.1016/j.tplants.2016.04.008.

40. Hao, L., Wang, H., Sunter, G., and Bisaro, D.M. (2003). Geminivirus AL2 and L2 proteins interact with and inactivate SNF1 kinase. Plant Cell 15, 1034–1048. 10.1105/tpc.009530.

41. Shen, W., Dallas, M.B., Goshe, M.B., and Hanley-Bowdoin, L. (2014). SnRK1 phosphorylation of AL2 delays Cabbage leaf curl virus infection in Arabidopsis. J Virol 88, 10598–10612. 10.1128/JVI.00761-14.

42. Schepetilnikov, M., Kobayashi, K., Geldreich, A., Caranta, C., Robaglia, C., Keller, M., and Ryabova, L.A. (2011). Viral factor TAV recruits TOR/S6K1 signalling to activate reinitiation after long ORF translation. EMBO J 30, 1343–1356. 10.1038/emboj.2011.39.

43. Filipe, O., De Vleesschauwer, D., Haeck, A., Demeestere, K., and Hofte, M. (2018). The energy sensor OsSnRK1a confers broad-spectrum disease resistance in rice. Sci Rep 8, 3864. 10.1038/s41598-018-22101-6.

44. De Vleesschauwer, D., Filipe, O., Hoffman, G., Seifi, H.S., Haeck, A., Canlas, P., Van Bockhaven, J., De Waele, E., Demeestere, K., Ronald, P., and Hofte, M. (2018). Target of rapamycin signaling orchestrates growth-defense trade-offs in plants. New Phytol 217, 305–319. 10.1111/nph.14785.

45. Meteignier, L.V., El Oirdi, M., Cohen, M., Barff, T., Matteau, D., Lucier, J.F., Rodrigue, S., Jacques, P.E., Yoshioka, K., and Moffett, P. (2017). Translatome analysis of an NB-LRR immune response identifies important contributors to plant immunity in Arabidopsis. J Exp Bot 68, 2333–2344. 10.1093/jxb/erx078.

46. Kalachova, T., Müller, K., Lacek, J., Pree, S., Antonova, A., Bondarenko, O., Burketová, L., Retzer, K., and Weckwerth, W. (2025). SnRK1.1 Coordinates Organ-Specific Growth–Defense Programs via Transcriptomic Rewiring in Arabidopsis thaliana. bioRxiv, 2025.2004.2025.650715. 10.1101/2025.04.25.650715.

47. Weigel, R.R., Bauscher, C., Pfitzner, A.J., and Pfitzner, U.M. (2001). NIMIN-1, NIMIN-2 and NIMIN-3, members of a novel family of proteins from Arabidopsis that interact with NPR1/NIM1, a key regulator of systemic acquired resistance in plants. Plant Mol Biol 46, 143–160. 10.1023/a:1010652620115.

48. Weigel, R.R., Pfitzner, U.M., and Gatz, C. (2005). Interaction of NIMIN1 with NPR1 modulates PR gene expression in Arabidopsis. Plant Cell 17, 1279–1291. 10.1105/tpc.104.027441.

49. Kinkema, M., Fan, W., and Dong, X. (2000). Nuclear localization of NPR1 is required for activation of PR gene expression. Plant Cell 12, 2339–2350. 10.1105/tpc.12.12.2339.

50. Zavaliev, R., Mohan, R., Chen, T., and Dong, X. (2020). Formation of NPR1 Condensates Promotes Cell Survival during the Plant Immune Response. Cell 182, 1093–1108 e1018. 10.1016/j.cell.2020.07.016.

51. Hrabak, E.M., Chan, C.W., Gribskov, M., Harper, J.F., Choi, J.H., Halford, N., Kudla, J., Luan, S., Nimmo, H.G., Sussman, M.R., et al. (2003). The Arabidopsis CDPK-SnRK superfamily of protein kinases. Plant Physiol 132, 666–680. 10.1104/pp.102.011999.

52. Kulik, A., Wawer, I., Krzywinska, E., Bucholc, M., and Dobrowolska, G. (2011). SnRK2 protein kinases--key regulators of plant response to abiotic stresses. OMICS 15, 859–872. 10.1089/omi.2011.0091.

53. Tang, R.J., Wang, C., Li, K., and Luan, S. (2020). The CBL-CIPK Calcium Signaling Network: Unified Paradigm from 20 Years of Discoveries. Trends Plant Sci 25, 604–617. 10.1016/j.tplants.2020.01.009.

54. Margalha, L., Valerio, C., and Baena-Gonzalez, E. (2016). Plant SnRK1 Kinases: Structure, Regulation, and Function. Exp Suppl 107, 403–438. 10.1007/978-3-319-43589-3_17.

55. Huber, S.C., and Huber, J.L. (1996). Role and Regulation of Sucrose-Phosphate Synthase in Higher Plants. Annu Rev Plant Physiol Plant Mol Biol 47, 431–444. 10.1146/annurev.arplant.47.1.431.

56. Sugden, C., Donaghy, P.G., Halford, N.G., and Hardie, D.G. (1999). Two SNF1-related protein kinases from spinach leaf phosphorylate and inactivate 3-hydroxy-3-methylglutaryl-coenzyme A reductase, nitrate reductase, and sucrose phosphate synthase in vitro. Plant Physiol 120, 257–274. 10.1104/pp.120.1.257.

57. Sugden, C., Crawford, R.M., Halford, N.G., and Hardie, D.G. (1999). Regulation of spinach SNF1-related (SnRK1) kinases by protein kinases and phosphatases is associated with phosphorylation of the T loop and is regulated by 5’-AMP. Plant J 19, 433–439. 10.1046/j.1365-313x.1999.00532.x.

58. Martinez-Barajas, E., and Coello, P. (2020). Review: How do SnRK1 protein kinases truly work? Plant Sci 291, 110330. 10.1016/j.plantsci.2019.110330.

59. Belda-Palazon, B., Adamo, M., Valerio, C., Ferreira, L.J., Confraria, A., Reis-Barata, D., Rodrigues, A., Meyer, C., Rodriguez, P.L., and Baena-Gonzalez, E. (2020). A dual function of SnRK2 kinases in the regulation of SnRK1 and plant growth. Nat Plants 6, 1345–1353. 10.1038/s41477-020-00778-w.

60. Peixoto, B., Moraes, T.A., Mengin, V., Margalha, L., Vicente, R., Feil, R., Hohne, M., Sousa, A.G.G., Lilue, J., Stitt, M., et al. (2021). Impact of the SnRK1 protein kinase on sucrose homeostasis and the transcriptome during the diel cycle. Plant Physiol 187, 1357–1373. 10.1093/plphys/kiab350.

61. Muralidhara, P., Weiste, C., Collani, S., Krischke, M., Kreisz, P., Draken, J., Feil, R., Mair, A., Teige, M., Muller, M.J., et al. (2021). Perturbations in plant energy homeostasis prime lateral root initiation via SnRK1-bZIP63-ARF19 signaling. Proc Natl Acad Sci U S A 118. 10.1073/pnas.2106961118.

62. Goransson, O., McBride, A., Hawley, S.A., Ross, F.A., Shpiro, N., Foretz, M., Viollet, B., Hardie, D.G., and Sakamoto, K. (2007). Mechanism of action of A-769662, a valuable tool for activation of AMP-activated protein kinase. J Biol Chem 282, 32549–32560. 10.1074/jbc.M706536200.

63. Hu, Y., Bai, J., Xia, Y., Lin, Y., Ma, L., Xu, X., Ding, Y., and Chen, L. (2022). Increasing SnRK1 activity with the AMPK activator A-769662 accelerates seed germination in rice. Plant Physiol Biochem 185, 155–166. 10.1016/j.plaphy.2022.06.005.

64. Kumar, S., Zavaliev, R., Wu, Q., Zhou, Y., Cheng, J., Dillard, L., Powers, J., Withers, J., Zhao, J., Guan, Z., et al. (2022). Structural basis of NPR1 in activating plant immunity. Nature 605, 561–566. 10.1038/s41586-022-04699-w.

65. Schmidt, E.K., Clavarino, G., Ceppi, M., and Pierre, P. (2009). SUnSET, a nonradioactive method to monitor protein synthesis. Nat Methods 6, 275–277. 10.1038/nmeth.1314.

66. Chen, T., Xu, G., Mou, R., Greene, G.H., Liu, L., Motley, J., and Dong, X. (2023). Global translational induction during NLR-mediated immunity in plants is dynamically regulated by CDC123, an ATP-sensitive protein. Cell Host Microbe 31, 334–342 e335. 10.1016/j.chom.2023.01.014.

67. Donaldson, S.G., Fox, O.F., Kishore, G.S., and Carubelli, R. (1981). Effect of manganese ions on the interaction between ribosomes and endoplasmic reticulum membranes isolated from rat liver. Biosci Rep 1, 727–731. 10.1007/BF01116471.

68. Mahfouz, M.M., Kim, S., Delauney, A.J., and Verma, D.P. (2006). Arabidopsis TARGET OF RAPAMYCIN interacts with RAPTOR, which regulates the activity of S6 kinase in response to osmotic stress signals. Plant Cell 18, 477–490. 10.1105/tpc.105.035931.

69. Van Leene, J., Han, C., Gadeyne, A., Eeckhout, D., Matthijs, C., Cannoot, B., De Winne, N., Persiau, G., Van De Slijke, E., Van de Cotte, B., et al. (2019). Capturing the phosphorylation and protein interaction landscape of the plant TOR kinase. Nat Plants 5, 316–327. 10.1038/s41477-019-0378-z.

70. Kang, S.A., Pacold, M.E., Cervantes, C.L., Lim, D., Lou, H.J., Ottina, K., Gray, N.S., Turk, B.E., Yaffe, M.B., and Sabatini, D.M. (2013). mTORC1 phosphorylation sites encode their sensitivity to starvation and rapamycin. Science 341, 1236566. 10.1126/science.1236566.

71. Din, F.V., Valanciute, A., Houde, V.P., Zibrova, D., Green, K.A., Sakamoto, K., Alessi, D.R., and Dunlop, M.G. (2012). Aspirin inhibits mTOR signaling, activates AMP-activated protein kinase, and induces autophagy in colorectal cancer cells. Gastroenterology 142, 1504–1515 e1503. 10.1053/j.gastro.2012.02.050.

72. Doshi, H., Spengler, K., Godbole, A., Gee, Y.S., Baell, J., Oakhill, J.S., Henke, A., and Heller, R. (2023). AMPK protects endothelial cells against HSV-1 replication via inhibition of mTORC1 and ACC1. Microbiol Spectr 11, e0041723. 10.1128/spectrum.00417-23.

73. Brunton, J., Steele, S., Ziehr, B., Moorman, N., and Kawula, T. (2013). Feeding uninvited guests: mTOR and AMPK set the table for intracellular pathogens. PLoS Pathog 9, e1003552. 10.1371/journal.ppat.1003552.

74. Kim, J., Kundu, M., Viollet, B., and Guan, K.L. (2011). AMPK and mTOR regulate autophagy through direct phosphorylation of Ulk1. Nat Cell Biol 13, 132–141. 10.1038/ncb2152.

75. Zeng, Y., Li, B., Huang, S., Li, H., Cao, W., Chen, Y., Liu, G., Li, Z., Yang, C., Feng, L., et al. (2023). The plant unique ESCRT component FREE1 regulates autophagosome closure. Nat Commun 14, 1768. 10.1038/s41467-023-37185-6.

76. Li, H., Liao, Y., Zheng, X., Zhuang, X., Gao, C., and Zhou, J. (2022). Shedding Light on the Role of Phosphorylation in Plant Autophagy. FEBS Lett 596, 2172–2185. 10.1002/1873-3468.14352.

77. Herzig, S., and Shaw, R.J. (2018). AMPK: guardian of metabolism and mitochondrial homeostasis. Nat Rev Mol Cell Biol 19, 121–135. 10.1038/nrm.2017.95.

78. Emanuelle, S., Hossain, M.I., Moller, I.E., Pedersen, H.L., van de Meene, A.M., Doblin, M.S., Koay, A., Oakhill, J.S., Scott, J.W., Willats, W.G., et al. (2015). SnRK1 from Arabidopsis thaliana is an atypical AMPK. Plant J 82, 183–192. 10.1111/tpj.12813.

79. Bergman, M.E., Evans, S.E., Kuai, X., Franks, A.E., Despres, C., and Phillips, M.A. (2023). Arabidopsis TGA256 Transcription Factors Suppress Salicylic-Acid-Induced Sucrose Starvation. Plants (Basel) 12. 10.3390/plants12183284.

80. Xie, Z., and Chen, Z. (1999). Salicylic acid induces rapid inhibition of mitochondrial electron transport and oxidative phosphorylation in tobacco cells. Plant Physiol 120, 217–226. 10.1104/pp.120.1.217.

81. Crozet, P., Margalha, L., Confraria, A., Rodrigues, A., Martinho, C., Adamo, M., Elias, C.A., and Baena-Gonzalez, E. (2014). Mechanisms of regulation of SNF1/AMPK/SnRK1 protein kinases. Front Plant Sci 5, 190. 10.3389/fpls.2014.00190.

82. Hedbacker, K., and Carlson, M. (2008). SNF1/AMPK pathways in yeast. Front Biosci 13, 2408–2420. 10.2741/2854.

83. Rawat, S.S., and Laxmi, A. (2025). Salicylic acid represses primary root growth through the Glucose-Target of Rapamycin-E2Fa pathway in Arabidopsis. bioRxiv, 2025.2001.2029.635412. 10.1101/2025.01.29.635412.

84. Xie, Z., Zhao, S., Li, Y., Deng, Y., Shi, Y., Chen, X., Li, Y., Li, H., Chen, C., Wang, X., et al. (2023). Phenolic acid-induced phase separation and translation inhibition mediate plant interspecific competition. Nat Plants 9, 1481–1499. 10.1038/s41477-023-01499-6.

85. Li, X., Chai, X., Lyu, H.N., Fu, C., Tang, H., Shi, Q., Wang, J., and Xu, C. (2023). Chemical proteomics reveals an ISR-like response elicited by salicylic acid in Arabidopsis. New Phytol 237, 1486–1489. 10.1111/nph.18646.

86. Silva, A.M., Wang, D., Komar, A.A., Castilho, B.A., and Williams, B.R. (2007). Salicylates trigger protein synthesis inhibition in a protein kinase R-like endoplasmic reticulum kinase-dependent manner. J Biol Chem 282, 10164–10171. 10.1074/jbc.M609996200.

87. Enganti, R., Cho, S.K., Toperzer, J.D., Urquidi-Camacho, R.A., Cakir, O.S., Ray, A.P., Abraham, P.E., Hettich, R.L., and von Arnim, A.G. (2017). Phosphorylation of Ribosomal Protein RPS6 Integrates Light Signals and Circadian Clock Signals. Front Plant Sci 8, 2210. 10.3389/fpls.2017.02210.

88. Reuber, T.L., and Ausubel, F.M. (1996). Isolation of Arabidopsis genes that differentiate between resistance responses mediated by the RPS2 and RPM1 disease resistance genes. The Plant cell 8, 241–249. 10.1105/tpc.8.2.241.

89. Dong, X., Mindrinos, M., Davis, K.R., and Ausubel, F.M. (1991). Induction of Arabidopsis defense genes by virulent and avirulent Pseudomonas syringae strains and by a cloned avirulence gene. Plant Cell 3, 61–72. 10.1105/tpc.3.1.61.

90. Bitrian, M., Roodbarkelari, F., Horvath, M., and Koncz, C. (2011). BAC-recombineering for studying plant gene regulation: developmental control and cellular localization of SnRK1 kinase subunits. Plant J 65, 829–842. 10.1111/j.1365-313X.2010.04462.x.

91. Gu, Y., Zebell, S.G., Liang, Z., Wang, S., Kang, B.H., and Dong, X. (2016). Nuclear Pore Permeabilization Is a Convergent Signaling Event in Effector-Triggered Immunity. Cell 166, 1526–1538 e1511. 10.1016/j.cell.2016.07.042.

92. Wang, W., Withers, J., Li, H., Zwack, P.J., Rusnac, D.V., Shi, H., Liu, L., Yan, S., Hinds, T.R., Guttman, M., et al. (2020). Structural basis of salicylic acid perception by Arabidopsis NPR proteins. Nature 586, 311–316. 10.1038/s41586-020-2596-y.

93. Schindelin, J., Arganda-Carreras, I., Frise, E., Kaynig, V., Longair, M., Pietzsch, T., Preibisch, S., Rueden, C., Saalfeld, S., Schmid, B., et al. (2012). Fiji: an open-source platform for biological-image analysis. Nat Methods 9, 676–682. 10.1038/nmeth.2019.

94. Labun, K., Montague, T.G., Krause, M., Torres Cleuren, Y.N., Tjeldnes, H., and Valen, E. (2019). CHOPCHOP v3: expanding the CRISPR web toolbox beyond genome editing. Nucleic Acids Res 47, W171–W174. 10.1093/nar/gkz365.

95. Ge, S.X., Jung, D., and Yao, R. (2020). ShinyGO: a graphical gene-set enrichment tool for animals and plants. Bioinformatics 36, 2628–2629. 10.1093/bioinformatics/btz931.

